# A double role of the Gal80 N-terminus in activation of transcription by Gal4p

**DOI:** 10.1101/2020.02.03.931568

**Authors:** Annekathrin Reinhardt-Tews, Rościsław Krutyhołowa, Christian Günzel, Sebastian Glatt, Karin D Breunig

## Abstract

The yeast galactose switch operated by the Gal4p-Gal80p-Gal3p regulatory module is a textbook model of transcription regulation in eukaryotes. The Gal80 protein inhibits Gal4p-mediated transcription activation by binding to the transcription activation domain. Inhibition is relieved by formation of an alternative Gal80-Gal3 complex. In yeasts lacking a Gal3p ortholog the Gal1 protein combines regulatory and enzymatic activity. The data presented here reveal a so-far unknown role of the Gal80 N-terminus in the mechanism of Gal4p activation. The N-terminus contains an NLS, which is responsible for nuclear accumulation of KlGal80p and galactokinase inhibition *in vitro.* Herein we propose a model where the N-terminus of KlGal80p reaches into the catalytic center of KlGal1p of the nuclear fraction of KlGal1p triggering dissociation of the KlGal80-KlGal4 complex. We corroborate this model by genetic analyses and structural modelling and provide a rationale for the divergent evolution of the mechanism activating Gal4p.

**Summary blurb:** Activation of gene expression by Gal4p in *K. lactis* requires an element in the N-terminus of KlGal80 that mediates nuclear import, KlGal1 interaction and galactokinase inhibition

## INTRODUCTION

Transcription regulation in response to the environment occurs in all organisms and is essential for cellular homeostasis. Studies on the response to changing nutrients in microorganisms have provided deep insight into the adaptive processes at the molecular level. Pioneering work on the regulation of transcription by galactose in yeast gave rise to the Douglas-Hawthorne model of the GAL regulon (Douglas and Hawthorne, 1972; Oshima, 1991). This model assumes that a gene specific transcription activator (Gal4p) is inhibited by Gal80p. When a yeast cell has access to galactose the Gal80p-mediated inhibition is relieved by the Gal3p protein, which functions as a galactose transducer.

These three regulatory proteins, namely Gal4p, Gal80p and Gal1/3p, represent the heart of the “galactose switch”. Gal4p transmits the information leading to high transcription rates to the transcription machinery. Gal80p mediates inhibition by binding with high affinity to a specific site overlapping the C-terminal activation domain of Gal4p (Keegan et al., 1986; Ma and Ptashne, 1987; Ansari et al., 1998). Gal1p, the enzyme that catalyzes galactose phosphorylation by ATP, senses the presence of galactose intracellularly by binding galactose and ATP directly, which increases the affinity between Gal1p and Gal80p. Therefore, insights into the formation of the Gal1-Gal80 complex are central to the understanding of the *GAL* switch and the activation of Gal4p. The above described trimeric regulatory module is present in *Kluyveromyces lactis* (Zenke et al., 1993; Zenke et al., 1996) whereas *Saccharomyces cerevisiae* harbors a more complex GAL switch (Johnston, 1987). As a result of a genome duplication event in the *Saccharomyces* lineage a paralog of Gal1p exists, named Gal3p. The Gal3p variant of Gal1p has lost enzymatic activity and seems to be dedicated to regulating Gal4p activation by formation of the Gal3-Gal80 complex (Hittinger and Carroll, 2007). The ScGal1p variant has only residual regulatory activity because its affinity to ScGal80p is lower than that of Gal3p (Lavy et al., 2016).

Gal4p has been shown to work as a transactivator, which is inhibited by Gal80p, in many types of higher eukaryotic cells (Fisher et al., 1988; Kakidani and Ptashne, 1988; Lin et al., 1988; Ma et al., 1988; Eliason et al., 2018) stressing the evolutionary conservation of the transcription apparatus. Multiple studies dissected in great detail how Gal4p and Gal80p regulate gene expression and the regulation of the *GAL* genes in yeast became a textbook model for eukaryotic gene regulation. Nevertheless, there are still many open questions concerning the molecular details of its regulation. Specifically, the mechanics of Gal4p activation is poorly understood.

In the *Kluyveromyces* lineage, which in contrast to *S. cerevisiae* has not undergone genome duplication, KlGal1p is a so-called “moonlightning” protein, which has two distinct functions, the enzymatic and the Gal3-like regulatory function. These functions can be separated by specific mutation (Meyer et al., 1991; Zenke et al., 1996, 1996; Vollenbroich et al., 1999; Hittinger and Carroll, 2007). Cross-complementation experiments have indicated and crystal structures of KlGal80p, ScGal80p as well as ScGal1p and ScGal3p have confirmed a high conservation of the Gal4p-Gal80p-Gal1/3 modules (Zenke et al., 1996; Menezes et al., 2003; Thoden et al., 2007; Thoden et al., 2008; Lavy et al., 2012). However, several differences have been documented. (i) *ScGAL3* is unable to provide the *Klgal1* regulatory function in *K. lactis* unless *KlGAL80* is replaced by *ScGAL80,* indicating some incompatibility between ScGal3p and KlGal80p (Zenke et al., 1996). (ii) KlGal80p is a nuclear protein whereas ScGal80p is nucleocytoplasmic (Anders et al., 2006). (iii) Species specific phosphorylation of Gal80p and Gal4p have been reported (Mylin et al., 1989; Mylin et al., 1990) but the role of post-translational modifications in Gal4p regulation is poorly understood. Therefore, it is not entirely clear if the mechanistic details of the GAL switch in *K. lactis* and *S. cerevisiae* are identical or not.

Here we have addressed species specificity by identifying the nuclear localization signal of KlGal80p. We show that the extreme N-terminus of the KlGal80p protein contains a nuclear localization signal (NLS) responsible for nuclear import of KlGal80p and at the same time it promotes formation of the KlGal80-KlGal1 complex. Both functions are overlapping and important for KlGal4p regulation in *K. lactis.* We propose a direct interaction between the KlGal80 N-terminus and the catalytic center of KlGal1p, which supports formation of a nuclear KlGal80-KlGal1 complex in *K. lactis.* In addition, we present a structural model of the complex based on the atomic structures of free KlGal80p and of the ScGal3-ScGal80 co-complex. Furthermore, we discuss the fact that despite significant sequence similarity the functions of the KlGal80p N-terminus are not conserved in *S. cerevisiae* in the light of the divergent evolution of the two yeast genera.

## RESULTS

### Nuclear localization of Gal80p in *K. lactis* depends on the N-terminus

The Gal80 protein of *K. lactis* (KlGal80p) differs in its subcellular distribution from its homolog in *S. cerevisiae*. Whereas ScGal80p is localized both, in nucleus and cytoplasm (nucleocytoplasmic), KlGal80p is exclusively nuclear (Anders, 2006; Anders et al., 2006). To identify the NLS, various sub-fragments of KlGal80p were fused to GFP and constitutively expressed in a *Klgal80* deletion mutant. Localization of the fusion proteins was analyzed by fluorescence microscopy (**Supplementary** Fig**. S1**). Besides full-length KlGal80p all protein segments conferring nuclear localization contained the N-terminus (**Supplementary** Fig. **S1**, **Fig. 1**). The smallest nuclear segment comprised only the first 39 amino acids (KlGal80-C1p, **Fig. 1B**). GFP-KlGal80p with a deletion of those N-terminal amino acids (**KlGal80-DC1p)** was found in the whole cell (**Fig. 1C**). Taken together these data clearly indicate that an NLS is located at the N-terminus of KlGal80p.

**Figure 1:**
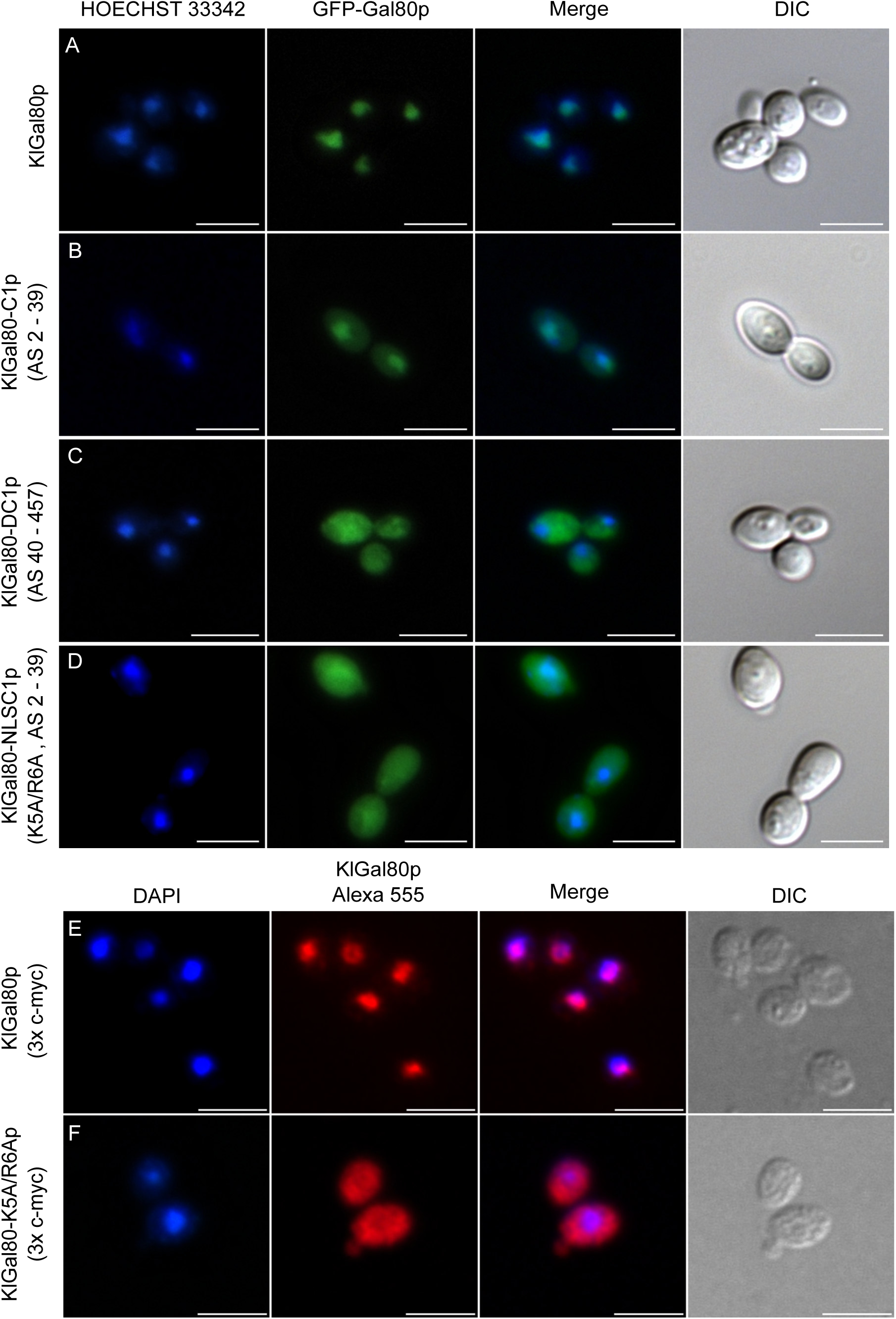
Identification of an N-terminal NLS in KlGal80p. The plasmids encoding GFP fused to (A) wild-type KlGal80p (pEG80WTGFPct), (B) the KlGal80 fragment aa 2-39 (pEQRS80C1), (C) the KlGal80 fragment aa 40 – 457 (pEQRS80DC1) and (D) the KlGal80 fragment aa 2-39 with exchange K5A/R6A (pEgal80NLS1C1) were transformed into the *Klgal80* deletion strain JA6/D802R. Cells were grown in minimal medium with 2 % glucose and analyzed by fluorescence microscopy. (E) Strains expressing c-myc-tagged wild-type KlGal80p (JA6/G80M) or (F) KlGal80-K5A/R6Ap (JA6/G80-KR56A) were grown in 2 % galactose and analyzed by immunofluorescence microscopy by a primary c-myc antibody and a secondary Alexa 555 antibody. The nucleus was stained by Hoechst 33342 or DAPI. Both channels were merged and a differential interference contrast (DIC) picture was taken. Scale bar: 5 µm.

Indeed, the KlGal80 N-terminus contains a sequence rich in basic residues ^5^KRSK^8^, which resembles a monopartite, class 2 NLS consensus sequence, K-K/R-X-K/R (Chelsky et al., 1989) (**Fig. 2, top**). To test the functionality of this sequence two of the basic residues, namely lysine 5 and arginine 6, were mutated (K5A R6A) by site-directed mutagenesis of the reading frame of GFP-KlGal80-C1p (aa 2-39). Like the N-terminal deletion, this point mutation abolished nuclear accumulation and showed nucleocytoplasmic localization (**Fig. 1D**). To rule out an effect of overexpression of the GFP-KlGal80-K5A/R6A protein, which was encoded by a multicopy plasmid, the mutation was also introduced into the chromosomal *KlGAL80* gene. Comparing the (“wild-type”) parent strain JA6/G80M encoding a (c-myc)3-tagged variant of KlGal80p with the resulting mutated strain JA6/G80-KR56A (*Klgal80-K5A/R6A*) by immunofluorescence microscopy confirmed that the basic residues K5 and R6 are important for nuclear localization. KlGal80-(c-myc)*3*p was nuclear under inducing conditions as shown before (Anders et al., 2006) whereas KlGal80-K5A/R6A-(c-myc)3p was nucleocytoplasmic (**Fig. 1E and 1F**).

**Figure 2:**
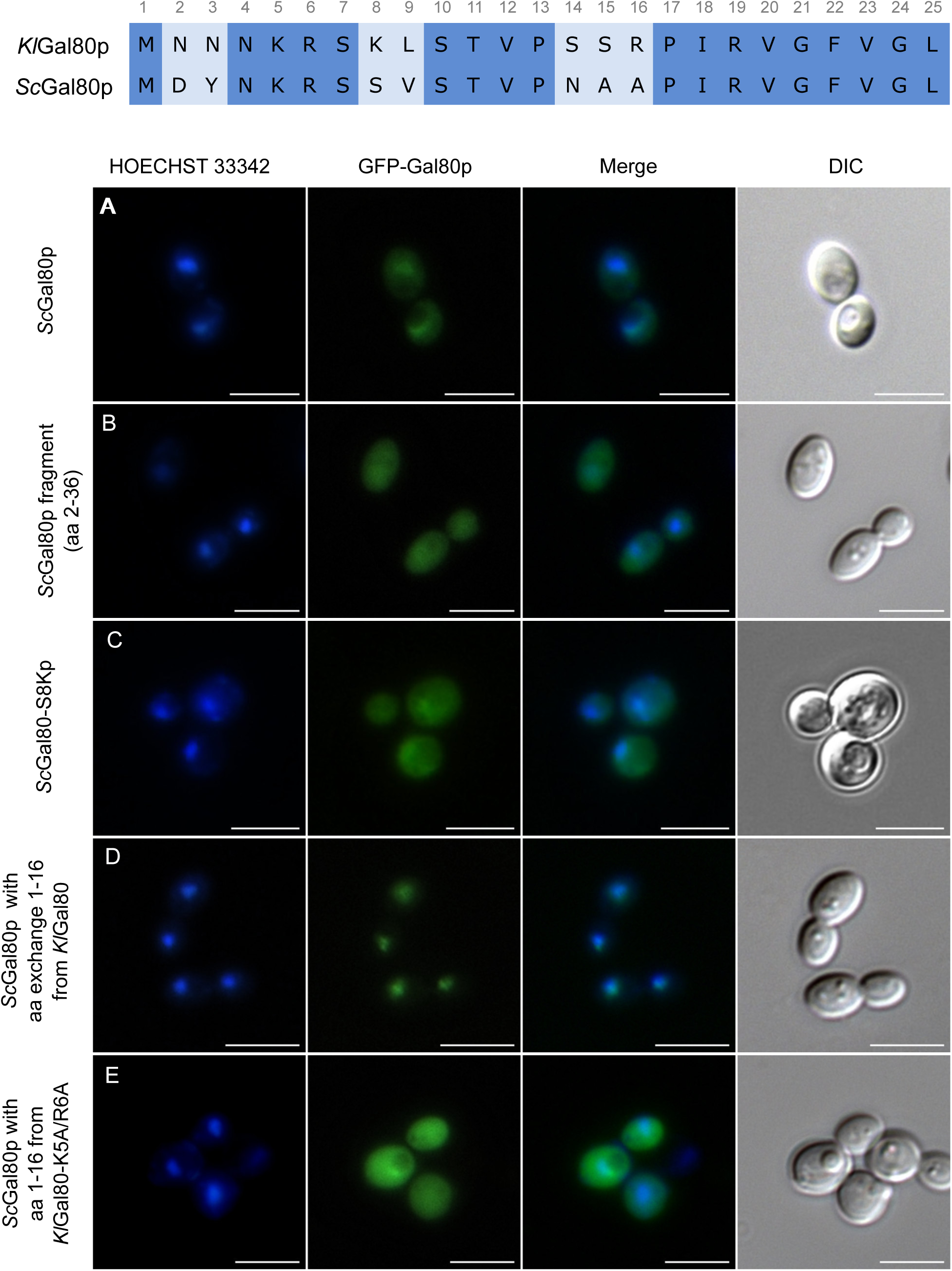
Comparative analysis of the N-termini of KlGal80p and ScGal80p. To analyze the function of the ScGal80p N-terminus plasmids encoding GFP fused ScGal80p-variants were transformed into the *K. lactis Klgal80* deletion strain JA6/D802R and localized by fluorescence microscopy. Cells expressing the plasmids (A) pEScG80 (wild-type ScGal80p), (B) pEScG8036 (ScGal80 fragment aa 2 - 36), (C) pEGFPScG80-S8K (ScGal80p with amino acid exchange S8K, (D) pEGFP-KlNT-ScG80 (ScGal80p with KlGal80p N-terminus aa 1-16) or (E) pEGFP-Kl56-ScG80 (ScGal80p with mutated KlGal80p N-terminus containing K5A/R6A exchange) were grown in minimal medium with 2 % glucose and analyzed by fluorescence microscopy. The nucleus was stained by Hoechst 33342. DIC: differential interference contrast. Scale bar: 5 µm.

### The 16 N-terminal amino acids determine the difference in subcellular distribution between ScGal80p and KlGal80p

It has been shown before that differences in subcellular distribution between KlGal80p and ScGal80p are an intrinsic property of the proteins and not of the host cell (Anders, 2006; Anders et al., 2006). ScGal80p in *K. lactis* is broadly distributed and KlGal80p in *S. cerevisiae* is nuclear, but both Gal80p homologues inhibit the heterologous Gal4p variant by interaction with the conserved Gal80p binding site in the C-terminal transcription activation domain (TAD). Here we show that the N-terminal 16 amino acids are responsible for the subcellular localization of the Gal80p variants (**Fig. 2**). A chimeric protein of ScGal80p with the N-terminus of KlGal80p (KlNTScGal80p) replacing the ScGAl80p N-terminus accumulated in the nucleus as efficiently as GFP-KlGal80p itself, whereas the mutated variant KlNTScGal80-K5A/R6Ap was also detected in the cytoplasm (**Fig. 2D, E**). This indicates (i) that ScGal80p does not have properties that prevent its nuclear accumulation and (ii) that these 16-aa are sufficient to cause nuclear accumulation of ScGal80p.

The N-termini of ScGal80p and KlGal80p are similar but deviate at seven positions within the first 16 amino acids (**Fig. 2** **top)**. ScGal80p (S8, A16) misses two basic residues (K8, R16), which are present in KlGal80p and partly constitute its NLS consensus motif ^5^KRSK^8^. To test whether the ScGal80p motif *^5^*KRSS*^8^* can be converted into an NLS the serine at position 8 was mutated to lysine giving the protein variant ScGal80-S8Kp. This protein, fused to GFP, was not exclusively nuclear but GFP fluorescence showed some nuclear accumulation (**Fig. 2C**)). Hence, the lysine at position 8 together with K5 and R6 contributes to nuclear transport in *K. lactis* but was not sufficient to create a strong NLS at the N-terminus of ScGal80p.

### Galactose induction is impaired in the *Klgal80-K5A/R6A* mutant

To address the question whether the reduced nuclear accumulation caused by the mutation in the KlGal80p-NLS had an influence on the KlGal80p inhibitor function KlGal4p controlled gene expression was examined. *LAC4*, one of the KlGal4p-regulated genes, encodes the *K. lactis* β-galactosidase, which allows easy monitoring of KlGal4p activity *in vivo*. Up to a certain limit the intensity of the blue color correlates very well with the *LAC4* transcript level, which in turn reflects the activity status of KlGal4p (Schaffrath and Breunig, 2000). Wild-type cells are white on glucose medium with X-Gal (because KlGal4p is inactive) and blue in the presence of an inducing sugar (lactose or galactose) due to KlGal4p activation and induction of the *LAC4* gene (**Fig. 3A**). A *Klgal80* deletion mutant is blue on both media because due to the absence of Gal80p KlGal4p is constitutively active. Both Gal80p variants, KlGal80p as well as ScGal80p, inhibit gene activation by binding to DNA-bound Gal4p in the nucleus. Hence, it was expected that the mutation in the KlGal80p NLS eventually might show weaker repression than wild-type KlGal80p due to lack of nuclear accumulation. To our surprise, the opposite was the case. The *K. lactis* strain JA6/G80-KR56A containing the *Klgal80-K5A/R6A* mutant allele was white also on galactose medium, indicating a super-repressed phenotype. Hence, the *Klgal80-K5A/R6A* mutant appears to be unable to respond to the presence of galactose.

**Figure 3:**
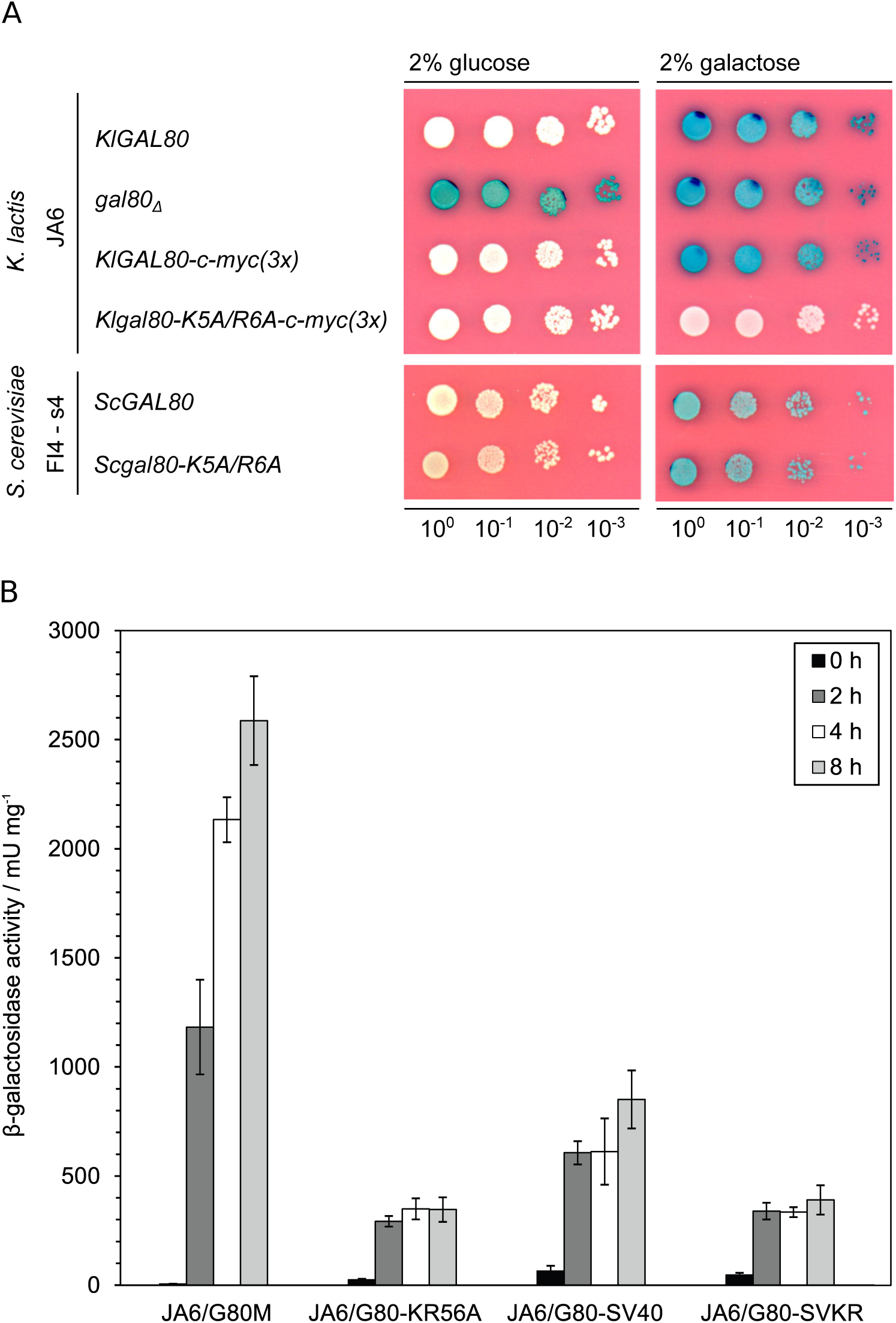
Gal4p controlled beta-galactosidase gene expression. Beta-galactosidase measurements to assay the influence of Gal80p localization on Gal4p controlled gene expression. **(A)** The *K. lactis* wild-type strain JA6 and derived mutated strains JA6/D802 (*Klgal80Δ*), JA6/G80M (c-myc-tagged *KlGAL80*) and JA6/G80-KR56A (c-myc-tagged *Klgal80-K5A/R6A*) and the *S. cerevisiae* strain FI4 s4 containing the *K. lactis LAC4-LAC12* genes and expressing wild-type ScGal80p or ScGal80-K5A/R6Ap were analyzed in an X-Gal plate assay. The plates were incubated for 3 days at 30 °C. **(B)** Strains expressing triple-myc-tagged wild-type Gal80p (JA6/G80M), Gal80-K5A/R6Ap (JA6/G80-KR56A) or the corresponding variants fused to SV40-NLS (JA6/G80-SV40) or (JA6/G80-SVKR) were grown in glucose and shifted to galactose containing medium. Beta-galactosidase activity was measured in crude extracts of samples taken at the indicated time points (hours) after the galactose shift. Mean values and standard variations based on two independent measurements each measuring three different dilutions with four technical replicates (12 replicates in total) are plotted.

β-galactosidase activity measurements over time in cells shifted from glucose to galactose medium (**Fig. 3B**) confirmed that in the wild-type cells there is an up to 700-fold induction of β-galactosidase activity. In contrast, the *Klgal80-K5A/R6A* mutant (strain JA6/G80-KR56A) showed only a weak short-term response (13-fold increase in 2 hrs) and no sustained induction over a period of at least 8 hours. To test the influence of the K5A/R6A mutation on ScGal80p function the K5A/R6A mutation was also introduced into the chromosomal *ScGAL80* gene of an *S. cerevisiae* strain. To have the same read-out as in *K. lactis* this strain carried the *LAC12* and *LAC4* genes of *K. lactis* and either the *ScGAL80* wild -type or the *Scgal80-K5A/R6A* mutant allele. On X-Gal containing plates both strains looked identical, they were white on glucose and blue on galactose medium (**Fig. 3A**, lower panel). Hence in contrast to *K. lactis*, the K5A/R6A mutation in ScGal80p did not affect Gal4p activation in response to galactose.

### The super-repressed phenotype is not caused by the altered localization of KlGal80-K5A/R6Ap

Next, we asked if in *K. lactis* the super-repressed phenotype of the *Klgal80-K5A/R6A* mutant is caused by the destruction of the NLS and thus to the resulting changes in intracellular localization of the mutated Gal80 protein. Therefore, we tried to redirect KlGal80-K5A/R6Ap to the nucleus by fusing the strong SV40-NLS (MGAPPKKKRKVA) to wild-type KlGal80p and to the mutated variant KlGal80-K5A/R6Ap, respectively. Immunofluorescence studies showed that the SV40-NLS did not interfere with the KlGal80p-NLS, as wild-type KlGal80p with and without the SV40-NLS was nuclear. Importantly, nuclear accumulation of mutant KlGal80-K5A/R6Ap could indeed be restored by the SV40-NLS (Supplementary Fig. S2D). We tested the ability of the KlGal80p variants to regulate KlGal4p controlled gene expression by measuring β-galactosidase activity after shifting from glucose to galactose (**Fig. 3B**). In *K. lactis* the fusion of KlGal80p with the heterologous SV40-NLS reduced galactose induction, but clearly did not prevent the response to galactose. The strains expressing the mutated KlGal80-K5A/R6Ap variant with or without the SV40-NLS only showed a weak, short-term response to the induction. Hence restoring the nuclear localization of the defective KlGal80-K5A/R6A protein did not restore inducibility, suggesting that the influence of the NLS-mutation on nuclear transport of KlGal80p is not enough to explain the super-repressed mutant phenotype.

### KlGal80p controls the intracellular distribution of KlGal1p

Since activation of Gal4p requires release of Gal80p from blocking the Gal4p-TAD and formation of the KlGal80-KlGal1 complex the simplest explanation for the non-inducible phenotype of the *Klgal80-K5A/R6A* mutant would be impaired in KlGal80-KlGal1 interaction.

The two amino acid substitutions might affect the stability of the complex directly or indirectly via the influence of KlGal80p on intracellular compartmentation.

Both in the presence and in the absence of KlGal80p, a GFP-KlGal1p fusion protein can be detected in the entire cell (Anders et al., 2006) (**Fig. 4A and B**). This changes with galactose induction or constitutive overexpression of the *KlGAL80* gene, which leads to nuclear accumulation of KlGal1p (**Fig. 4C**) (Anders et al., 2006). Apparently, the KlGal1p subcellular distribution is influenced by the concentration of KlGal80p and perhaps by the affinity between KlGal80p and KlGal1p.

**Figure 4:**
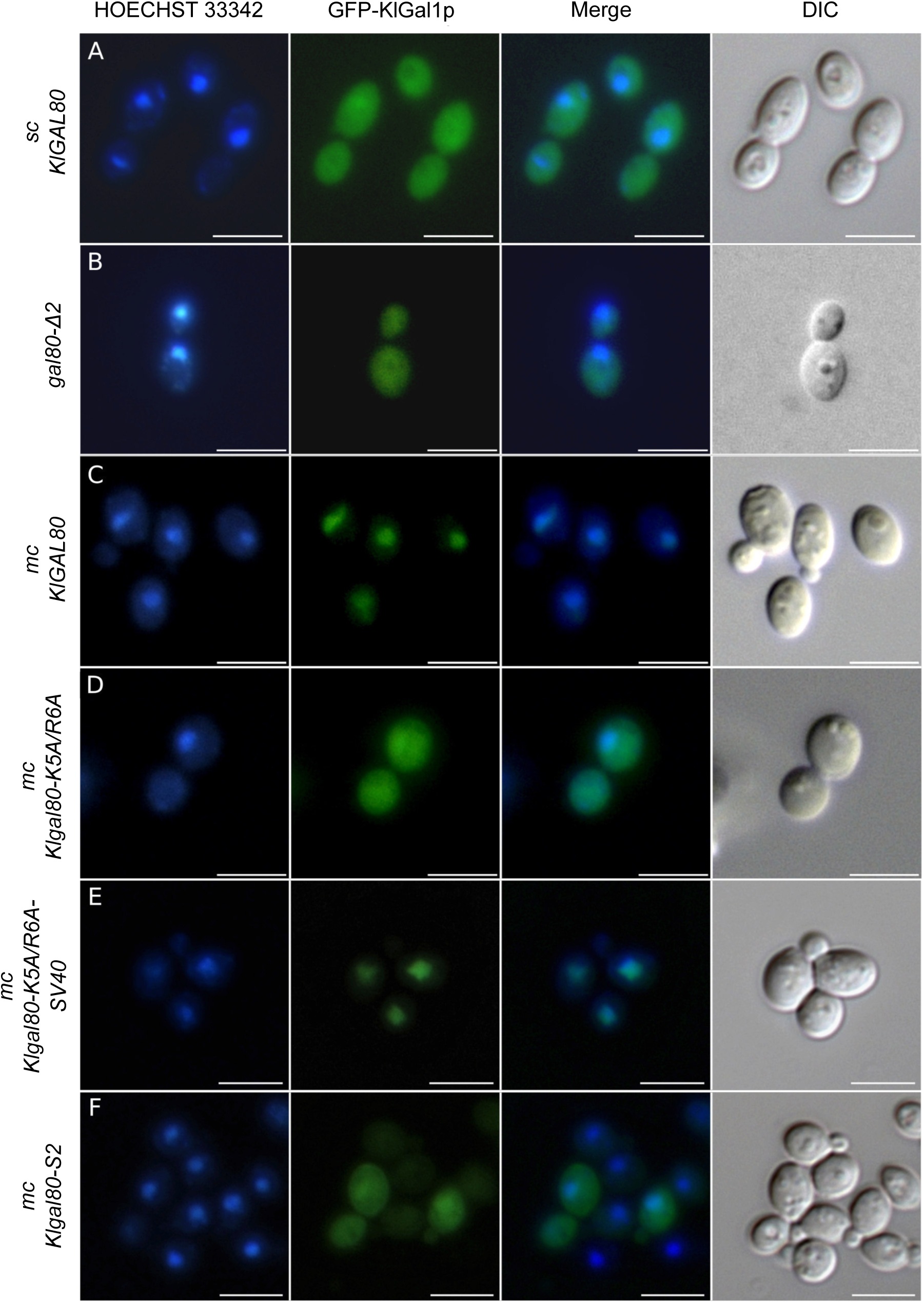
Influence of *Klgal80* mutations on KlGal1p subcellular localization. The KlGal1-GFP fusion protein was localized by fluorescence microscopy in cells expressing KlGal80p variants. The KlGAL1-GFP protein was constitutively expressed from a centromeric plasmid pCGFPAG1 (**A-B**) or pCGFPAG1-ura3Δ (**C-F**), the KlGal80p variants from a multicopy plasmid, both genes under control of the constitutive *ScADH1* promoter in *K. lactis* strain JA6/D1R (**A**) deleted for the chromosomal *KlGAL1* gene, or JA6/D1D802R deleted for *KlGAL1* and *KlGAL80* (**B-E**). KlGal80p variants were wild-type (expressed from chromosomal *KlGAL80* gene) (**A**); wild-type from multicopy plasmid pEAG80 (**C**); Gal80-K5A/R6Ap (from pEAG80-KR56A) (**D**); Gal80-K5A/R6Ap with N-terminal SV40-NLS-fusion (from pEAG80-KR56A-SV40) (**E**) or KlGal80^s2^p (E367K) from pEAG80S2 (**F**). The nucleus was stained with Hoechst 33342, KlGal1p was detected by its GFP-tag. sc - single copy. mc - multicopy. Merge: GFP and Hoechst channels were superimposed. DIC: differential interference contrast. Scale bar: 5 µm.

Here we addressed the question whether the mutation in the NLS of KlGal80p also impacts on the intracellular distribution of KlGal1p. When the mutated *Klgal80-K5A/R6A* allele was introduced in multicopy into a *Klgal80 Klgal1* mutant constitutively expressing the GFP-KlGal1p fusion protein no nuclear accumulation was observed (**Fig. 4D**). Since KlGal80-K5A/R6Ap itself does not accumulate in the nucleus (**Fig.1F**), we tested whether the SV40-NLS fusion, which partially restored nuclear accumulation of the KlGal80p-K5A/R6Ap variant, was able to promote nuclear accumulation of KlGal1p when overexpressed. This could indeed be shown (**Fig. 4E**). We also assayed a *K. lactis* mutant, *Klgal80-S2,* that is KlGal1p binding deficient due to an E367K exchange (Zenke et al., 1996). The KlGal80-E367K protein has an intact N-terminal NLS and is nuclear (Supplementary Fig. S3E). In contrast to nuclear KlGal80-K5A/R6A-SV40p, it did not accumulate GFP-KlGal1p in the nucleus (**Fig. 4F**). We conclude that accumulation of KlGal1p requires effective nuclear transport of KlGal80p, which can be achieved by the intact N-terminal NLS, or by a heterologous NLS, and (ii) the ability of KlGal1p and KlGal80p to form a complex. In summary, we propose that KlGal1p enters the nucleus piggy-backing on KlGal80p.

### The non-inducible phenotype of the *Klgal80-K5A/R6A* mutant is suppressed by overexpression of KlGal1p

To further elucidate why the NLS mutant is non-inducible we analyzed the influence of *KlGAL80* and *KlGAL1* gene dosage on KlGal4p activity using the X-Gal plate assay as in Fig. 3 (**Fig. 5**). The *K. lactis* strain JA6 (*KlGAL80)* (**Fig. 5A**) and an otherwise isogenic strain JA6/KR56A (*Klgal80*-*K5A/R6A*) (**Fig. 5B**) were transformed with single copy (sc) or multicopy (mc) plasmids containing the *KlGAL80* or the *KlGAL1* genes. As expected, in the *KlGAL80* wild-type background galactose induction is evident by the color shift from almost white on 2 % glucose to dark blue on galactose (**Fig. 5A**, empty vector). The unexpected blue color on 0,2 % glucose was partially suppressed by one additional gene copy (scGAL80) and fully suppressed by multicopy *KlGAL80* (mcGAL80). This correlation with the *KlGAL80* gene dosage indicates incomplete KlGal4p inhibition by KlGal80p under these growth conditions in the empty vector control. Strikingly, in the *Klgal80-K5A/R6A* mutant background the blue staining on low glucose plates was very similar to that of the wild-type indicating a low influence of the NLS-mutation in low glucose. On galactose plates, as expected, all transformants of the wild-type strain were dark blue confirming effective activation of KlGal4p whereas transformants of the mutated strain varied considerably, both in coloring and growth on low galactose medium (**Fig. 5B**). Most importantly, in the mutant background the non-inducible and the slow growth phenotypes were strongly suppressed by multicopy *GAL1* (mc GAL1). An additional *KlGAL1* gene copy (scGAL1) had a similar but weaker effect.

**Figure 5:**
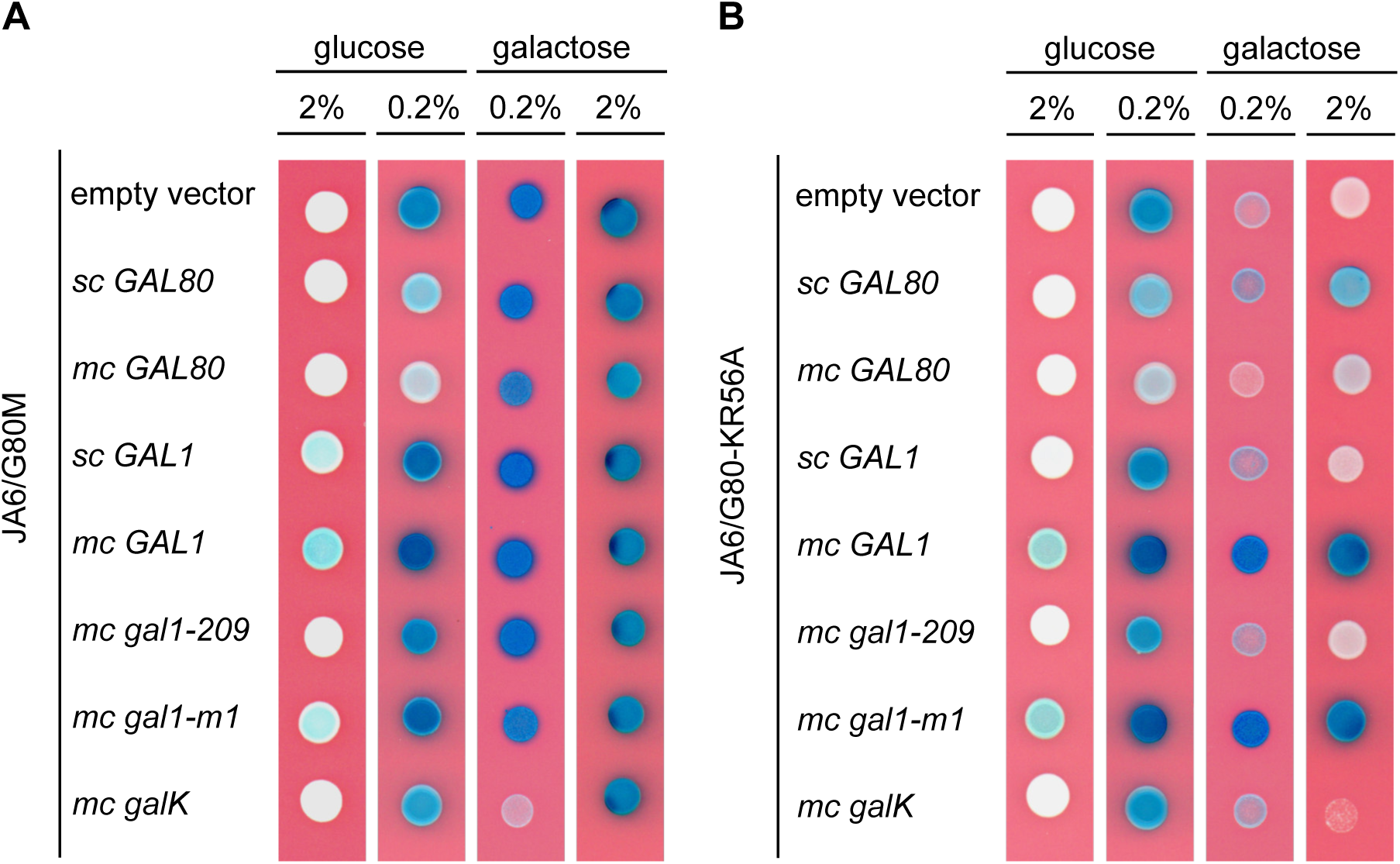
Influence of the *KlGAL1* gene on Gal4p controlled beta-galactosidase gene expression in *KlGAL80* wild-type and *Klgal80-K5A/R6A* mutant. The *K. lactis* strains **(A)** JA6/G80M (c-myc-tagged *KlGAL80*) and **(B)** JA6/G80-KR56A (c-myc-tagged *Klgal80-K5A/R6A*) were transformed with the multicopy (pE) or centromeric (pC) plasmids pE1 (empty vector), pC80 (*sc KlGAL80*), pEAG80 (*mc KlGAL80*), pCAG1 (*sc KlGAL1*), pEAG1 (*mc KlGAL1*), pEAG1-209 (mc *Klgal1-209*, N261Y), pEAG1-m1 (*mc Klgal1-m1*, E254K) and pEAGK1 (*mc galK* galactokinase from *E. coli*). The resulting colonies were analyzed in an X-Gal plate assay on YNB plates without uracil. The plates were incubated for 3 days at 30 °C. sc – single copy. mc – multicopy.

The suppressing effect of KlGal1p overexpression was not observed with the *Klgal1-209* allele in multicopy but was also seen with the *Klgal1-m1* allele. In KlGal1-209p an N261Y exchange causes loss of enzymatic activity but this protein can still bind to KlGal80p and promote galactose induction (Zenke et al., 1996; 1999). *Klgal1-m1* carries the E254K mutation that interferes with KlGal80 complex formation *in vitro* but is enzymatically active (Zenke et al., 1996; Zenke et al., 1999; Menezes et al., 2003). Since these phenotypes suggested that, unexpectedly, galactokinase activity might be more important for suppression than the ability to bind KlGal80p, we also tested whether the galactokinase gene from *E. coli* (*GalK)* in multicopy could suppress the *Klgal80-K5A/R6A* mutant phenotype but this was not the case.

As mentioned above overexpression of KlGal80p in the *Klgal80-K5A/R6A* mutant background revealed that, in contrast to other super-repressed Sc*GAL80^s^* alleles (Nogi and Fukasawa, 1984) the mutation is not dominant over the wild-type. Single copy *KlGAL80* (scGAL80) weakly improved induction by galactose and growth on galactose but the negative influence on KlGal4p activation of KlGal80p overexpression (Zenke et al., 1993) nullified this effect (**Fig. 5B**).

In summary, the finding that the non-inducible phenotype of the *Klgal80-K5A/R6A* mutant was efficiently suppressed by overexpression of KlGal1p or by a single copy of wild-type *KlGAL80* supports the view that KlGal1p and KlGal4p compete for binding to KlGal80p. However, the phenotypes associated with the two mutated KlGal1p variants cannot be easily explained by a simple equilibrium shift model (See also Discussion Section).

### The K5A/R6A mutation in KlGal80p affects KlGal1p interaction

We expressed and purified N-terminally His6-tagged versions of wild-type KlGal80p (NHKlGal80p), the K5A/R6A variant (NHKlGal80-K5A/R6Ap) and NHKlGal1p in bacteria to compare the affinities of wild-type and mutated KlGal80p for KlGal1p more directly. As described before, the interaction of KlGal80p with KlGal1p inhibits the galactokinase activity of this enzyme and can be used to quantify the interaction between the two proteins (Anders et al., 2006). Titration of increasing amounts of KlGal80p into a galactokinase assay resulted in a first order inhibition curve which enabled determination of dissociation constants (**Fig. 6A and B**). WT KlGal80p tightly binds KlGal1p (K_D_ ∼80nM) while the NHKlGal80-K5A/R6Ap indicates almost 40-fold weaker binding (K_D_ ∼3.1μM).

**Figure 6:**
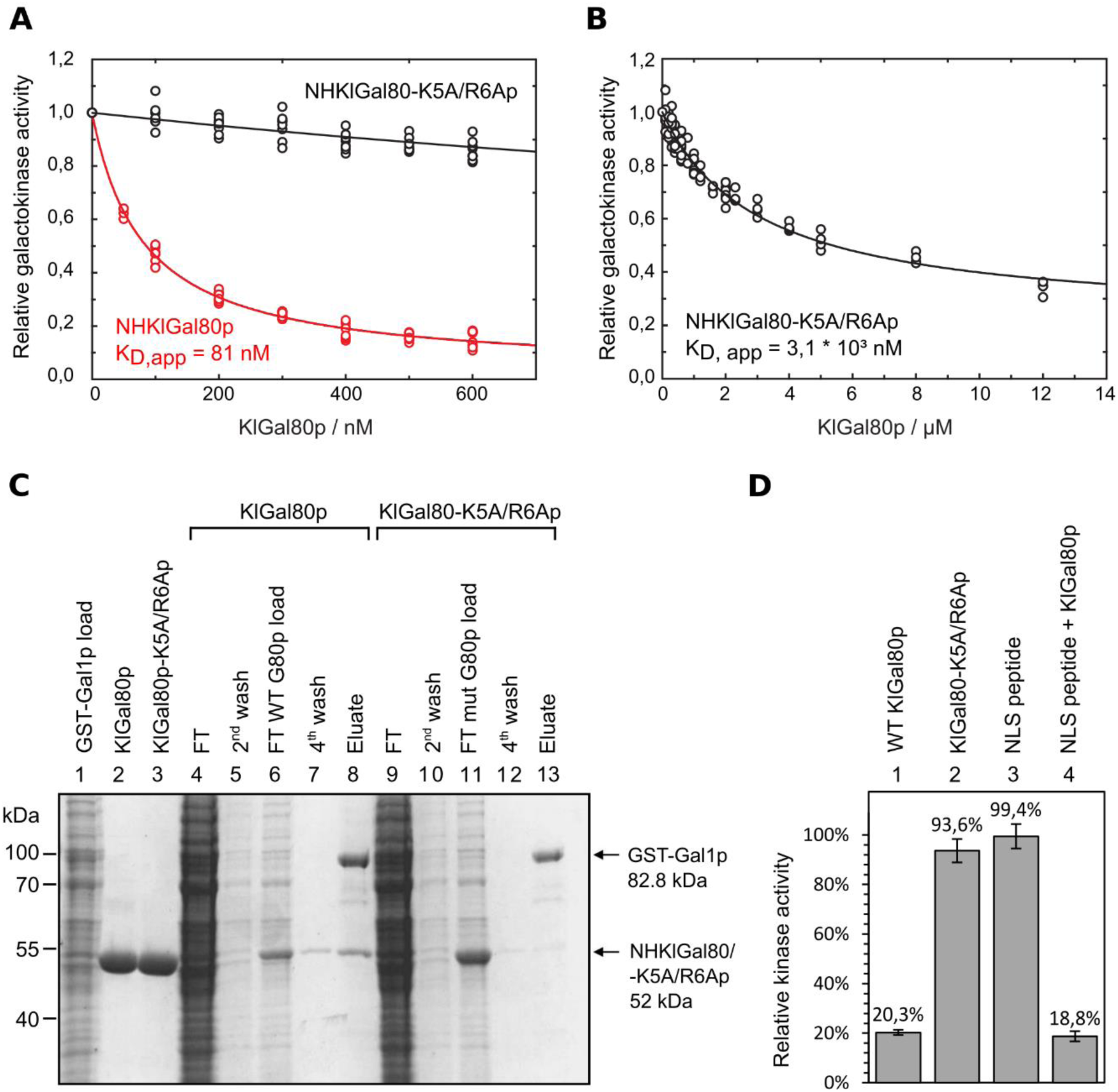
Influence of the KlGal1-NLS mutation on *Kl*Gal1p interaction. **(A** and **B)** Recombinant proteins produced in *E. coli* (KlGal80p with N-terminal His6-tag, NHKlGal80p, NHKlGal80-K5A/R6Ap and Gst-KlGa1p) were purified to determine the affinity of KlGal80p and KlGal80K5A/R6Ap for KlGal1p *in vitro* by making use of the inhibitory effect of KlGal80p on KlGal1p’s galactokinase activity. Enzymatic activity was determined as described previously (Anders et al., 2006). Data points result from two independent measurements with four replicates each. K_D, app_ values were calculated from the decline in KlGal1p activity with increasing KlGal80p (red, NHKlGal80p) or KlGal80-K5A/R6Ap (black, NHKlGal80-K5A/R6Ap) concentrations and were. K_D, app_ = 81 nM for KlGal80p wild-type and K_D, app_ = 3<u>.1</u> * 10^3^ nM for KlGal80-K5A/R6Ap. **(C)** The interaction of purified wild-type (IHGal80p) and mutant (IHGal80-K5A/R6Ap) KlGal80p with GST-tagged KlGal1p was analyzed by a GST-Pull-Down-Assay. 1500 µg raw extract containing GST-Gal1p was immobilized on a GST-column for 1.5 hour and then washed twice. 20 µg of purified Gal80p was added and incubated for 1.5 hour. The column was washed four times and GST-Gal1p was eluted with buffer containing 40 mM glutathion. 165 µg of raw extract from JA6/D1D802R transformed with pGSTGal1, 2 µg of purified KlGal80p as well as 10 µl of flow through, washing and elution fractions were loaded on an SDS-Gel and stained with Coomassie solution after PAGE. WF – wash fraction, FT – flow through. (**D**) Comparing the inhibition of galactokinase activity of full-length KlGal80p, KlGal80-K5A/R6Ap and a synthetic 17 aa NLS peptide. The influence is expressed as relative galactokinase activity in the presence vs absence of the Gal80 variant. Apparently, the peptide alone has neither negative nor positive influence.

A lower affinity of the NHKlGal80-K5A/R6Ap to KlGal1p compared to wild-type was also confirmed in a GST pull-down assay (**Fig. 6C**). Whereas wild-type KlGal80p could be detected in the glutathione elution fraction containing GST-KlGal1p indicating elution of the complex, this was not the case for KlGal80-K5A/R6Ap. Apparently, only the wild-type KlGal80p could be trapped in complex with GST-Gal1p on the glutathione matrix. Therefore, we conclude that the N-terminal NLS-mutation results in a lower affinity for KlGal1p *in vitro*. The inhibitory activity of purified KlGal80p-K5R/R6Ap vs. KlGal80p is plotted in (**Figure 6D**). To test whether the N-terminus of KlGal80p alone can bind KlGal1p, we used a synthetic peptide corresponding to amino acid 1-17 of KlGal80p. The presence or absence of that peptide did not show any difference in KlGal80p mediated galactokinase inhibition *in vitro*, suggesting that the N-terminus is essential, but not sufficient for the interaction with KlGal1p. Taken together these findings support the view that the super-repressed phenotype of the NLS mutant is caused by a weaker interaction between KlGal80-K5A/R6Ap and KlGal1p.

### N-terminus of KlGal80p can bind near the active site of KlGal1p

So far sequence conservation, point mutations that affect galactose induction and the structure of the ScGal80-ScGal3 complex (Vollenbroich et al., 1999; Menezes et al., 2003; Lavy et al., 2012) had shown that direct contacts between Gal3p and Gal80p reside mostly in the C-terminal half of both proteins. But upon formation of the KlGal1-KlGal80 complex the N-terminus of KlGal80p could be placed in close vicinity of the KlGal1p active site. Therefore, we decided to analyze preferential orientations of the KlGal80p N-terminus on the KlGal1p-KlGal80 complex *in silico*. Based on a surface electrostatic charge of a modelled KlGal1-KlGal80 complex (Supplementary Fig. S4A) five potential binding sites were defined (Supplementary Fig. S4 B). Subsequently, peptide docking of KlGal80(1-16) wild-type to these potential binding sites resulted in a model with the highest likelihood of binding (**Fig. 7A**). In the energetically most favorable conformation the N-terminus would clash with the bound ATP molecule (**Fig. 7B**), potentially locking phosphogalactose in the catalytic center of Gal1p. In the proposed model the interaction of KlGal80(1-16) wild-type with KlGal1p is mostly dependent on N3, K5 and R6. This could explain the differences in the rate of inhibition of galactokinase activity by KlGal80p wild-type and KlGal80-K5A/R6Ap (**Fig. 6A** **and** B). When docking of the mutated N-terminus of KlGal80(1-16)-K5A/R6A to the KlGal1-KlGal80 complex was performed the active site of KlGal1p was occupied with a lower probability (Supplementary Fig. S4C, D). The predictions are in line with the obtained enzyme kinetic data (**Fig. 6 A, B**) and the decreased affinity of the KlGal80-K5A/R6Ap to KlGal1p (**Fig. 6C**). Based on our biochemical and bioinformatical studies, we would like to propose that in addition to an extended Gal1p-Gal80p interface involving the KlGal80p C-terminus there is an additional contact site between the KlGal80p-N-terminus and KlGal1p. This latter contact may initiate the crucial step in KlGal4p activation leading to KlGal4p release and induction of *GAL/LAC* genes expression (**Fig. 7C**).

**Figure 7:**
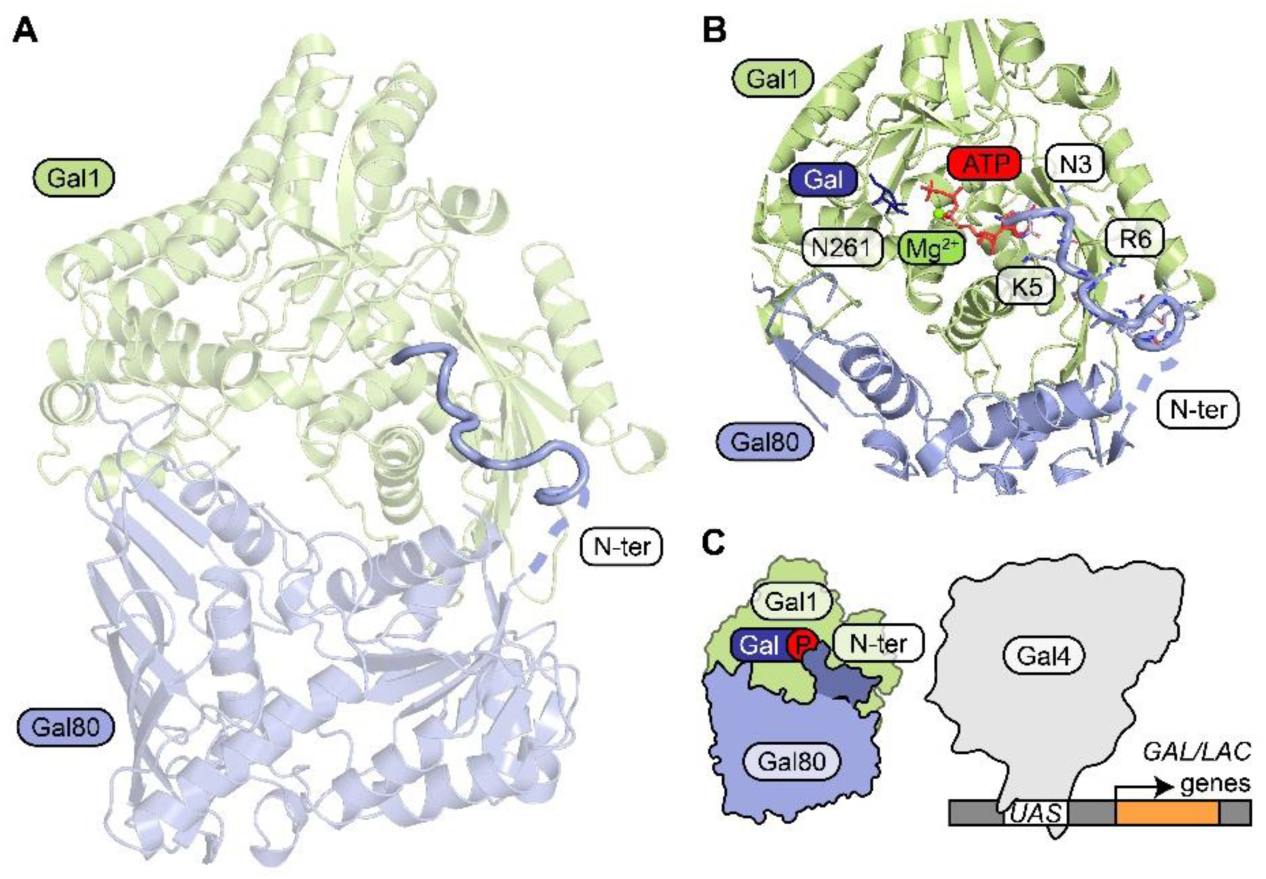
Structural model of KlGal80p-KlGal1p-interaction and the role of the KlGal80 N-terminus. **(A)** Structural overview of a homology-based model of KlGal1 (based on ScGal3 in 3V2U) in complex with KlGal80 (PDB ID 3E1K). Thick cartoon (N-terminus, blue) represents the best docking result of KlGal80 1-16 to the KlGal1 molecule. Only one heterodimer of KlGal1-KlGal80 complex is shown. **(B)** A close-up of KlGal1p active site in an open conformation. Galactose (dark blue), magnesium (bright green) and ATP (red) were positioned according to their position in ScGal3 (PDB ID 3V2U). Docked KlGal80 1-16 peptide clashes with ATP. Residues of potential importance for peptide-protein interactions are highlighted. **(C)** A schematic representation of KlGal1-KlGal80 complex. The N-terminus of KlGal80 (darker blue) blocks the ATP binding site of KlGal1p, effectively locking phosphorylated galactose inside. Gal1-Gal80 interaction leads to the release of KlGal4p, which activates galactose metabolism-related gene expression.

## DISCUSSION

The *Saccharomyces* and *Kluyveromyces* genomes have been shaped by more than 100 million years of divergent evolution. In that time two genetic events occurred that undoubtedly had massive impact on the GAL regulon. Firstly, the whole genome duplication in the *Saccharomyces* lineage was accompanied by rearrangements, deletions and duplications of chromosome segments (Lynch and Conery, 2000; Kellis et al., 2004) and secondly the loss of the *LAC* genes disabled lactose metabolism, which occurred in *S. cerevisiae* and independently in many other ascomycetes. The structures and functions of the regulatory proteins involved in the GAL switch are very similar in *S. cerevisiae* and *K. lactis* but there is also evidence for divergent evolution at the protein level. Here we report data supporting the view that in detail the molecular mechanics of Gal4p activation is not identical in the two yeast species.

We have asked how the reported differences in intracellular distribution between KlGal80p and ScGal80p (Anders et al., 2006) are achieved and maintained. We could show that the very N-terminus of KlGal80p contains an autonomous NLS able to direct a GFP-fusion protein to the nucleus. A class 2 NLS consensus ^5^K-K/R-X-K/R^8^ is a necessary functional element of the KlGal80-NLS. It was sufficient to swap the 16 N-terminal amino acids of ScGal80p and KlGal80p to exchange their intracellular distribution giving ScGal80p an exclusive nuclear KlGal80p-like localization and *vice versa.* Unexpectedly, changing the localization of KlGal80p from exclusively nuclear to nucleocytoplasmic by mutating the basic amino acids K5 and R6 in the NLS impaired KlGal4p activation. Since the super-repressed phenotype was not affected by redirecting KlGal80-K5A/R6Ap to the nucleus we concluded that impaired activation of KlGal4p in the mutant is not due to impaired nuclear import of KlGal80-K5A/R6Ap but to a so-far unknown property of the N-terminus of KlGal80p. It is shown here that the N-terminal region involving the NLS consensus sequence is not only important for nuclear transport of KlGal80p but also for protein-protein interaction with KlGal1p and inhibition of KlGal1p’s galactokinase activity. These functions are overlapping and are all affected by the introduced NLS-mutation. The negative influence of the mutation on Gal4p activated gene expression identified the N-terminus of *K. lactis* Gal80p as an important new element in KlGal4p activation.

The fact that the super-repressed phenotype of the NLS mutant can be suppressed by overexpression of KlGal1p is compatible with the competition model for Gal4p activation where Gal4p and Gal1/3p compete for the binding to Gal80p (Bhat and Hopper, 1992; Anders, 2006; Jiang et al., 2009). But equilibrium binding constants and on- and off rates for the Gal1/Gal3-Gal80 and Gal4-Gal80 complexes measured *in vitro* (Verma et al., 2004; Anders and Breunig, 2010; Abramczyk et al., 2012; Pannala et al., 2012; Lavy et al., 2016) indicated that the transition of Gal80p from one complex to the other cannot be explained by a genuine equilibrium shift. All models of the Gal4p activation mechanism have in common that a change in Gal80p and/or Gal3p is assumed that substantially lowers the binding constant of the Gal80-Gal4 complex (Wightman et al., 2008; Egriboz et al., 2013).

We propose a model where the interaction of the KlGal80p N-terminus with nuclear KlGal1p triggers a structural change in KlGal80p, which promotes dissociation of the KlGal4-Gal80 complex. According to the model, the KlGal80p variant with the mutated N-terminus would be impaired in triggering this change as it is impaired in inhibiting the galactokinase activity.

In principle, the structurally ill-defined N-terminus of KlGal80p can reach out to the catalytic center of the galactokinase enzyme. In our *in silico* docking experiments with the modeled KlGal1-KlGal80 complex and a KlGal80p 1-16 N-terminal peptide the preferred docking site on KlGal1p for the N-terminus of KlGal80p was oriented towards the bound ATP whereas with the mutated peptide this preference was not seen. Our model implies that in addition to the conserved major interface of the Gal80-Gal1/Gal3 complex (Lavy et al., 2012) there is a second site of interaction in the KlGal80-KlGal1 complex which seems to be conserved only in the genus *Kluyveromyces*.

The bifunctionality of KlGal1p provides a rational for compartmentation of the regulators. We have shown that accumulation of KlGal80p recruits KlGal1p to the nucleus in a way involving the N-terminal NLS. Since interaction between KlGal80p and KlGal1p results in inhibition of the catalytic activity of KlGal1p (Anders et al., 2006), formation of galactose-1-phosphate (Gal-1P), the first step in galactose metabolism, would be slowed down as KlGal80p accumulates. However, the overlap of the KlGal80-NLS with the region causing galactokinase inhibition provides a means to restrict enzyme inhibition to the nuclear fraction of KlGal1p. In the cytoplasm binding of the NLS to an importin might occlude the N-terminal KlGal1p interaction site but should allow the co-transport of KlGal1p and KlGal80p via the major interaction site. In the nucleus, where dissociation of the importin would occur, the NLS would be released and could find a docking site on KlGal1p preferentially reaching into the substrate containing catalytic center. This would stabilize the KlGal1-KlGal80 complex in a ligand bound manner and trigger a conformational change in KlGal80p (Vollenbroich et al., 1999) that lowers the affinity of the KlGal80-KlGal4 complex leading to activation of KlGal4p.

This model makes a number of assumptions that can be tested experimentally. It postulates a transient interaction of all three components of the regulatory module as proposed for *S. cerevisiae* Gal4p, Gal80p and Gal3p (Abramczyk et al., 2012). After all, the mechanism of Gal4p activation in *S. cerevisiae* may not be entirely different from that in *K. lactis.* There is evidence that ScGal4p also requires ScGal3p and ScGal1p to enter the nucleus (Egriboz et al., 2011), that a region at the N-terminus of ScGal80p is involved in nuclear transport (Nogi and Fukasawa, 1989) and, last but not least, that a trimeric complex precedes the dissociation of Gal4p and Gal80p. However, the coupling of KlGal1p nuclear transport to that of KlGal80p seems to be unique and an elegant manifestation of optimization in the simpler setting of the regulatory module in *Kluyveromyces spp*.

## MATERIAL AND METHODS

### Yeast Strains and Growth Conditions

The *K. lactis* strains (Supplementary table S1) JA6/D802 (*gal80-Δ2::ScURA3*) (Zenke et al., 1993), JA6/D1R (*gal1::Scura3*) (Zachariae, 1994), JA6/D1D802R (*gal80-Δ2::Scura3 gal1::Scura3*) (Zachariae, 1994), JA6/G80M (*GAL80*), JA6/G80-KR56A (*gal80-K5A, R6A*), JA6/G80-SV40 (*SV40-NLS-GAL80*), JA6/G80-SVKR (*SV40-NLS-gal80-K5A, R6A*), were congenic to JA6 (*MAT α ade1-600 adeT-600 ura3-12 trp1-11*) (Breunig and Kuger, 1987) and differed in the indicated alleles. The SV40-NLS (MGAPPKKKRKVA) was fused N-terminally. JA6/G80M, JA6/G80-KR56A, JA6/G80-SV40 and JA6/G80-SVKR contained a C-terminal c-Myc-epitope (3x). The different *KlGAL80* alleles were generated by site-directed mutagenesis using the QuikChange Multi Site-Directed Mutagenesis Kit (Agilent Technologies) or by fusion PCR, verified by DNA sequencing and introduced into JA6/D802 (*Klgal80::ScURA3*) (Zenke et al., 1993) as restriction fragments replacing *Klgal80::ScURA3*.

The *S. cerevisiae* strains FI4-s4 ScGAL80 (*ScGAL80*) and FI4-s4 ScGAL80KR56A (*Scgal80-K5A, R6A*) were isogenic to FI4 sin4ΔScgal80Δ (*sin4::HIS5 Scgal80::ScURA3*) (Schmidt, D., PhD thesis, 2010) except for the indicated *ScGAL80* alleles. The different *GAL80* alleles were generated by site-directed mutagenesis using the QuikChange Multi Site-Directed Mutagenesis Kit (Agilent Technologies) or by fusion PCR, verified by DNA sequencing and introduced into FI4 sin4ΔScgal80Δ as a PCR fragment replacing *Scgal80::ScURA3*. Yeast cells were grown in synthetic minimal medium (0.67 % yeast nitrogen base supplied with an amino acid/base mix). Glucose and galactose were added as carbon sources with a final concentration of 2 %. For selection of prototroph transformants, the corresponding amino acid or base was left out.

### Plasmids and Cloning

The multicopy vector pEG80WTGFPct codes for a C-terminally GFP-fused KlGal80p expressed under control of the ScADH1 promoter (Supplementary table S2). The PCR fragment obtained with KlG80SmiIBw (Supplementary table S3) and MluATGWTG80fw from pEgal80NLS1GFPct was cut with *Mlu*I and *Smi*I and ligated with the equally cut vector pEgal80NLS1GFPct. The multicopy plasmids pEQRS80A1 - C5 code for GFP-fused KlGal80p fragments expressed under control of the ScADH1 promoter. The PCR fragments obtained with the corresponding primers (Supplementary table S4) from pEQRS80 (Hager, 2003) were cut with *Mlu*I and *Smi*I and ligated with the equally cut pEQRS80 vector. The multicopy vector pEQRS80DC1 codes for a GFP-fused KlGal80 fragment (residue 40-457) expressed under control of the ScADH1 promoter. The PCR fragment obtained with C2MluIKlG80FwNeu and KlG80SmiIBw from pEQRS80 was cut with *Mlu*I and *Smi*I and ligated with the equally cut vector pEQRS80. The multicopy vector pEgal80NLS1C1 codes for a GFP-fused KlGal80p fragment (residues 2-39) with the amino acid exchange K5A/R6A expressed under control of the ScADH1 promoter. The PCR fragment obtained with GFPFw and C1SmiIKlG80Bw from pEgal80NLS1 was cut with *Mlu*I and *Smi*I and ligated with the equally cut vector pEQRS80. The multicopy vector pEScG80 codes for a GFP-fused ScGal80p expressed under control of the ScADH1 promoter. The PCR fragment obtained with MluIScGal80fw and SwaIScGal80bw from pScGal80 was cut with *Mlu*I and *Smi*I and ligated with the equally cut vector pEQRS80. The multicopy vector pEScG8036 codes for a GFP-fused KlGal80p fragment (residues 2-36) expressed under control of the ScADH1 promoter. The PCR fragment obtained with GFPFw and ScG80_36_Smi_Bw from pEScG80 was cut with *Mlu*I and *Smi*I and ligated with the equally cut vector pEQRS80. The multicopy vector pEGFP-KlNT-ScG80 codes for a GFP-fused ScGal80p variant, where the N-terminal 16 amino acids were exchanged with those of KlGal80p, expressed under control of the ScADH1 promoter. The PCR fragment obtained with KlG80_rv and KlG80NT_15AS_fw from pEScG80 was cut with *Mlu*I and *Smi*I and ligated with the equally cut vector pEQRS80. The multicopy vector pEGFP-Kl56-ScG80 codes for a GFP-fused ScGal80p variant, where the N-terminal 16 amino acids were exchanged with those of KlGal80-K5A/R6Ap, expressed under control of the ScADH1 promoter. The PCR fragment obtained with KlG80_rv and KlG80_KR56A_NT15AS_fw from pEScG80 was cut with *Mlu*I and *Smi*I and ligated with the equally cut vector pEQRS80. pGSTGal1 codes for a glutathione S-transferase-KlGal1 fusion protein (Zenke et al., 1999). The centromeric plasmid pCGFPAG1 codes for a GFP-KlGAL1 fusion protein expressed under control of the ScADH1 promoter (Anders et al., 2006). The multicopy vector pEGFPScG80-S8K codes for a GFP-fused ScGal80p variant with the amino acid exchange S8K expressed under control of the ScADH1 promoter. The PCR fragment obtained with KlG80_rv and ScG80_K8_fw from pEScG80 was cut with *Mlu*I and *Smi*I and ligated with the equally cut vector pEScG80. The Plasmid pCGFPAG1-ura3Δ was obtained by introducing a frameshift mutation into the ScURA3 gene of pCGFPAG1 (Anders et al., 2006). The centromeric plasmids pCGal1HA and pCNLSGal1HA code for C-terminally HA-tagged KlGal1 without (pCGal1HA) and with an N-terminal SV40-NLS (MGAPPKKKRKVA) (pCNLSGal1HA). The multicopy plasmid pEAG80S2 codes for the variant KlGal80-S2p (E367K) expressed under control of the ScADH1 promoter (Zenke et al., 1999). The multicopy plasmid pEAG80KR56A codes for the *Klgal80* double mutant K5A, R6A under control of the ScADH1 promoter. The *Klgal80-KR56A* containing *Bcu*I and *Pfl23II* fragment of pAG80KR56A was cloned into pEAG80 (Zenke, 1995). pAG80KR56A was constructed by site directed mutagenesis using the primers KlG80KR56A_fw and Amp_DS with pAG80 (Zenke et al., 1999). The plasmid pEAG80-KR56A-SV40 codes for a KlGal80p variant with the exchange K5A/R6A and an N-terminal SV40-NLS-fusion (MGAPPKKKRKVA). pEAG80-KR56A-SV40 was constructed by InFusion cloning of the *Smi*I and *Spe*I cut vector pEAG80 and the PCR products obtained with the primers ADH1-G80-inFu-FW and ADH1-G80-inFu-RV from pEAG80 as well as G80-KR56A-InFu-FW and G80-KR56A-inFu-RV from pI80KR56ASV40. The integration plasmid pI80KR56ASV40 codes for KlGal80p with the exchange K5A/R6A, C-terminal c-myc-tag (3x) and an N-terminal SV40-NLS-fusion (MGAPPKKKRKVA) and was obtained by integration of the *Eco*81I and *Van*91I cut fusion PCR fragment into the same cut vector pI80Myc. The fusion PCR was performed with the primers Integration_pIG80_3 and KlGAL80-422C and the PCR fragments of SV40NLSKlG80_bw and Integration_pIG80_3 as well as SV40NLS_KR56A_fw and KlGAL80-422C with pI80KR56A as the template. The integration plasmid pI80KR56A codes for KlGal80p with the amino acid exchange K5A/R6A and was constructed by ligation of the pKlGal80KR56A *Van*91I and *Eco*81I fragment containing the upstream fragment of *Klgal80-K5A/R6A* with the equally cut pI80Myc. The plasmid pKlGal80KR56A coding for KlGal80p with the exchange K5A/R6A was constructed by site directed mutagenesis with the QuikChange multi site-directed mutagenesis kit (Agilent Technologies) with the primers KlGal80NLS1CO and Amp_DS and the plasmid pKlGal80 (Zenke et al., 1993). The integration plasmid pI80Myc coding for c-myc-tagged (3x) KlGal80p was constructed with the QuikChange Multisite-Directed Mutagenesis Kit, a megaprimer containing the c-myc-tag and the plasmid pI80 (Zenke et al., 1993). The megaprimer was obtained by PCR with the primers TagFw and MycTagBw on pYM5 (Knop et al., 1999). Plasmid pETNHG80KR56A is a derivative of pETNHG80 (Anders et al., 2006) and codes for the N-terminal His6-tagged KlGal80p variant (K5A, R6A, position in the untagged protein). Plasmid pETIHG80KR56A is a derivative of pETIHG80 (Anders et al., 2006) and codes for the internal His6-tagged KlGal80p variant (K5A, R6A).

### Protein Expression and Purification

N-terminally or internal 6xHis-tagged wild-type KlGal80p (NHKlGal80p or IHGal80p), double mutant KlGal80-K5A/R6Ap (NHKlGal80-K5A/R6Ap or IHGal80-K5A/R6Ap) and KlGal1p (NHGal1p) were expressed in Escherichia coli strain Rosetta (DE3)-pLys from pET-derived (Novagen) expression vectors (pETNHG80, pETIHG80, pETNHG1 (Anders et al., 2006), pETNHG80-KR56A and pETIHG80KR56A. Purification was executed as previously described (Anders et al., 2006).

### Peptide Synthesis

The peptide KlN17G80 included the N-terminal 17 residues of KlGal80p (MNNNKRSKLSTVPSSRP) and was synthesised by the group of Prof. Dr. Frank Bordusa, Martin-Luther-University, Halle, Germany.

### Galactokinase inhibition assay

The galactokinase inhibition experiments were performed as described previously (Anders et al., 2006). Shortly ADP production from ATP in the reaction (galactose + ATP → galactose − 1 − phosphate + ADP) was coupled to the reactions of pyruvate kinase and lactate dehydrogenase monitoring NADH consumption. Each reaction mixture contained 10 nM NHGal1p and increasing concentrations of NHKlGal80p or NHKlGal80-KR56Ap. Samples containing the different KlGal80p variants were measured against a sample with NHKlGal1p alone giving relative galactokinase activities. K_D, app_ values were calculated from the decline in KlGal1p activity with increasing KlGal80p concentrations.

### Immunofluorescence microscopy

Immunofluorescence slides were prepared according to the protocol of Audrey L. Atkin (Atkin, 1998). Nuclear staining was performed with DAPI or Hoechst 33342. Primary antibodies were c-Myc (monoclonal, from mouse, by Roche) and c-Myc (polyclonal, from rabbit, by Santa Cruz). Secondary antibody was Alexa Fluor 555 (anti rabbit, life technologies). Spheroblasting of cells was done with Zymolyase 100T (Seikagaku). Cells expressing GFP-fused proteins were resuspended in PBS containing 4 % formaldehyde and dropped on polyline coated slides. After 5 minutes the slides were washed with PBS and the cells were coated with 10 µl Hoechst 33342 (5 µg/ml) and 10 µl Mounting Medium (according to Atkin, 1998) and coverslips.

### X-Gal plate assay

A cell suspension of a single colony was diluted and spotted on YNB agar plates containing the indicated carbon source, an amino acid / base mix and X-Gal. Plates were incubated at 30 °C and documented by scanning after three days.

### Preparation of yeast protein extracts

Logarithmically grown cells were resuspended in extraction buffer (50 mM HEPES-KOH pH 7,3; 60 mM potassium acetate; 5 mM magnesium acetate; 0,05 % TritonX 100 v/v; 10 % glycerol w/v; 1 mM sodium fluoride; 20 mM glycerophosphate; 1 mM DTT; complete protease inhibitor (Roche)) and disrupted by vortexing together with glass beads.

### β-Galactosidase assay

β-galactosidase activity of protein extracts was measured by detecting the increase in light absorbance at 420 nm in buffer containing 5 mM Tris/HCl pH 7,8; 5 % (v/v) glycerol, 10 mM KCl and 4 mg/ml o-nitrophenyl-β-D-galactopyranoside (ONPG) at 30°C in a 96 well plate with a SpectrostarNano plate reader.

### GST-pull-down assay

Protein raw extracts were isolated from a logarithmically grown culture of JA6/D1D802R transformed with pGSTGal1 by vortexing with glass beads in lysis buffer containing 50 mM Tris/HCl pH 7,5, 150 mM NaCl, 10 % glycerol, 0,1 % Tween 20 and 1,4 mM 2-mercaptoethanol. 1500 µg protein extract containing GST-Gal1p was loaded on a GST SpinTrap column (GE Healthcare) and incubated for 1.5 hour at 4°C. The column was washed twice with 600 µl binding buffer (lysis buffer + 2 % galactose and 5 mM ATP). 20 µg IHGal80/-K5A/R6A in binding buffer were incubated on the column for 1.5 hour rotating at 4°C. The column was washed three times with 600 µl and once with 200 µl binding buffer. The proteins were eluted with 200 µl elution buffer (binding buffer containing 40 mM glutathion).

### Structural modelling and docking

Homology-based model of KlGal1p was prepared using SWISS-MODEL server (Waterhouse et al., 2018) with ScGal3 (PDB ID 3V2U) as a template. The obtained model of KlGal1p was combined with the known KlGal80p structure (PDB ID 3E1K) and positioned according to ScGal3p-ScGal80p position in PDB ID 3V2U. The protein-protein interface of the modelled complex was validated using InterProSurf (Negi et al., 2007). PyMOL software was used for visualization purposes (DeLano). Surface charge distributions were calculated using APBS electrostatics (Jurrus et al., 2018). The KlGal80p 1-16 peptide was placed in five arbitrarily chosen areas of the KlGal1p-KlGal80p model based on surface charge and presence of a potential binding cleft. Prepared models for WT KlGal80 1-16 and K5A/R6A mutant peptide were submitted to FlexPepDock server (Raveh et al., 2010; London et al., 2011). Movements within the KlGal1-KlGal80 complex were restrained while the KlGal80p 1-16 peptide could undergo up to 10 Å of movement. 100 low resolution models and 100 high resolution models were prepared for of each position. The ten models with lowest Rosetta scores (which indicates highest binding probability) were collected.

## ACKNOWLEDGEMENTS

We thank Katharina Böhm for construction of the pEGFPScG80-S8K plasmid. This work was supported by grant BR921/7 and BR921/9-1 (KDB and ART). In addition, RK and SG were supported by grant OPUS16 2018/31/B/NZ1/03559 from the Polish National Science Centre. We thank the MCB structural biology core facility (supported by the TEAM TECH CORE FACILITY/2017-4/6 grant from Foundation for Polish Science) for providing computational resources.

## AUTHOR CONTRIBUTIONS

ART performed all yeast, fluorescence, biochemical, biophysical and activity assays; CG contributed experimental data shown in Figure 6C and D. RK performed structural modeling with input from SG. KDB envisioned the project, designed the experimental concepts and, together with ART, prepared the first manuscript draft. All authors contributed to the final text of the manuscript and to the ART, RK, SG and KDB preparation of figures.

## CONFLICT OF INTERESTS

The authors declare no competing interests.

---

**Supplementary Figure S1:**
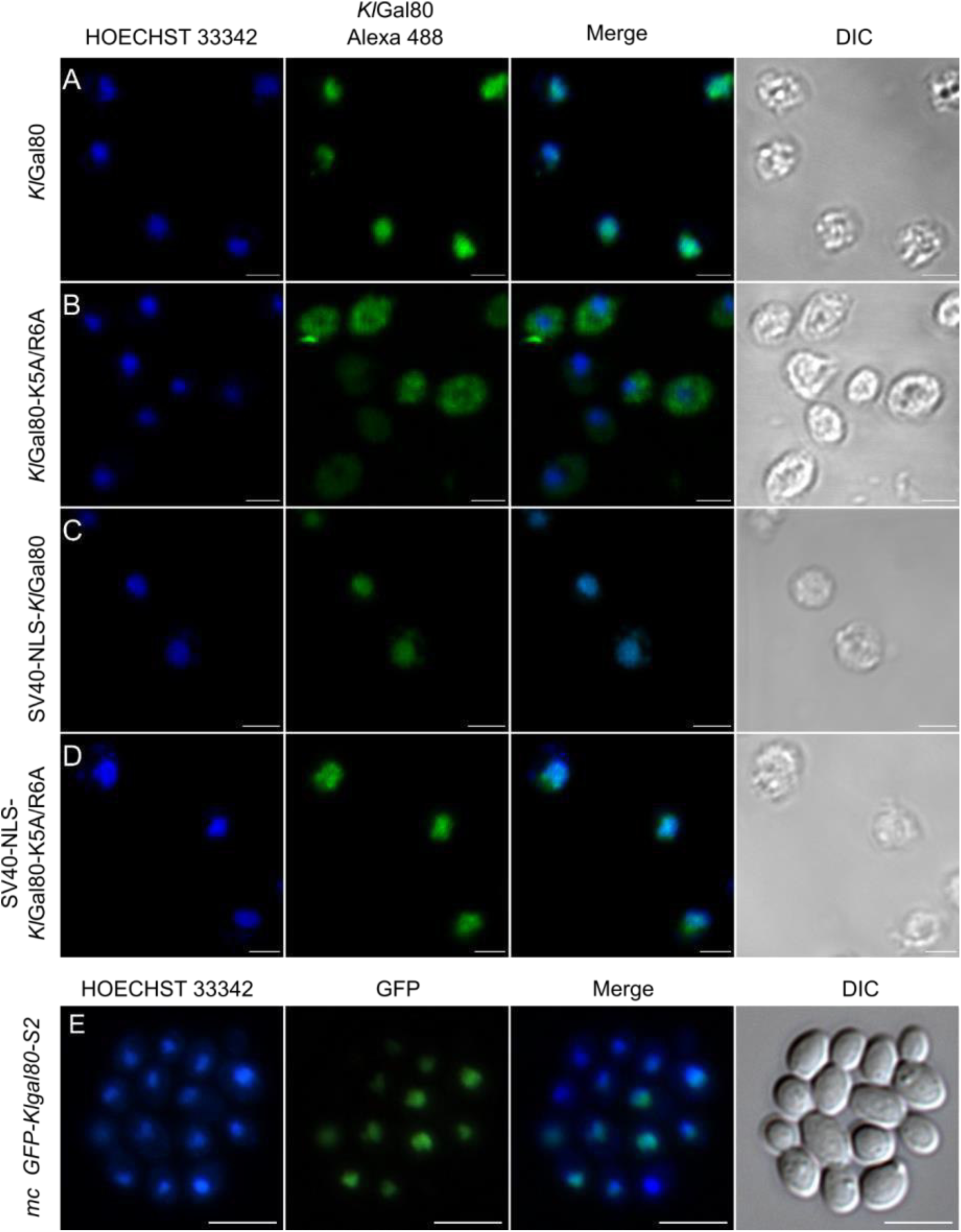
Mapping of nuclear localization signal in KlGal80p by and fluorescence microscopy of GFP-fusion proteins. Top: Overview of the PCR-generated segments of KlGal80p fused to the N-terminus of GFP. Nuclear localization is indicated by a bold frame. Bottom: the plasmids (**A**) pEQRS80-A1, (**B**) pEQRS80-A2, (**C**) pEQRS80-A3 and (**D**) pEQRS80-B1, (**E**) pEQRS80-B2, (**F**) pEQRS80-B3,(**G**) pEQRS80-C1, (**H**) pEQRS80-C2, (**I**) pEQRS80-C3, (**J**) pEQRS80-C4 and (**K**) pEQRS80-C5 coding for GFP fused KlGal80-fragments were transformed into the *Klgal80* deletion strain JA6/D802R. Cells were grown in minimal medium with 2 % glucose and analyzed by fluorescence microscopy. The nucleus was stained by Hoechst 33342. The channels for GFP and Hoechst were merged and a differential interference contrast (DIC) picture was taken. Scale bar: 5 µm.

**Supplementary Figure S2:**
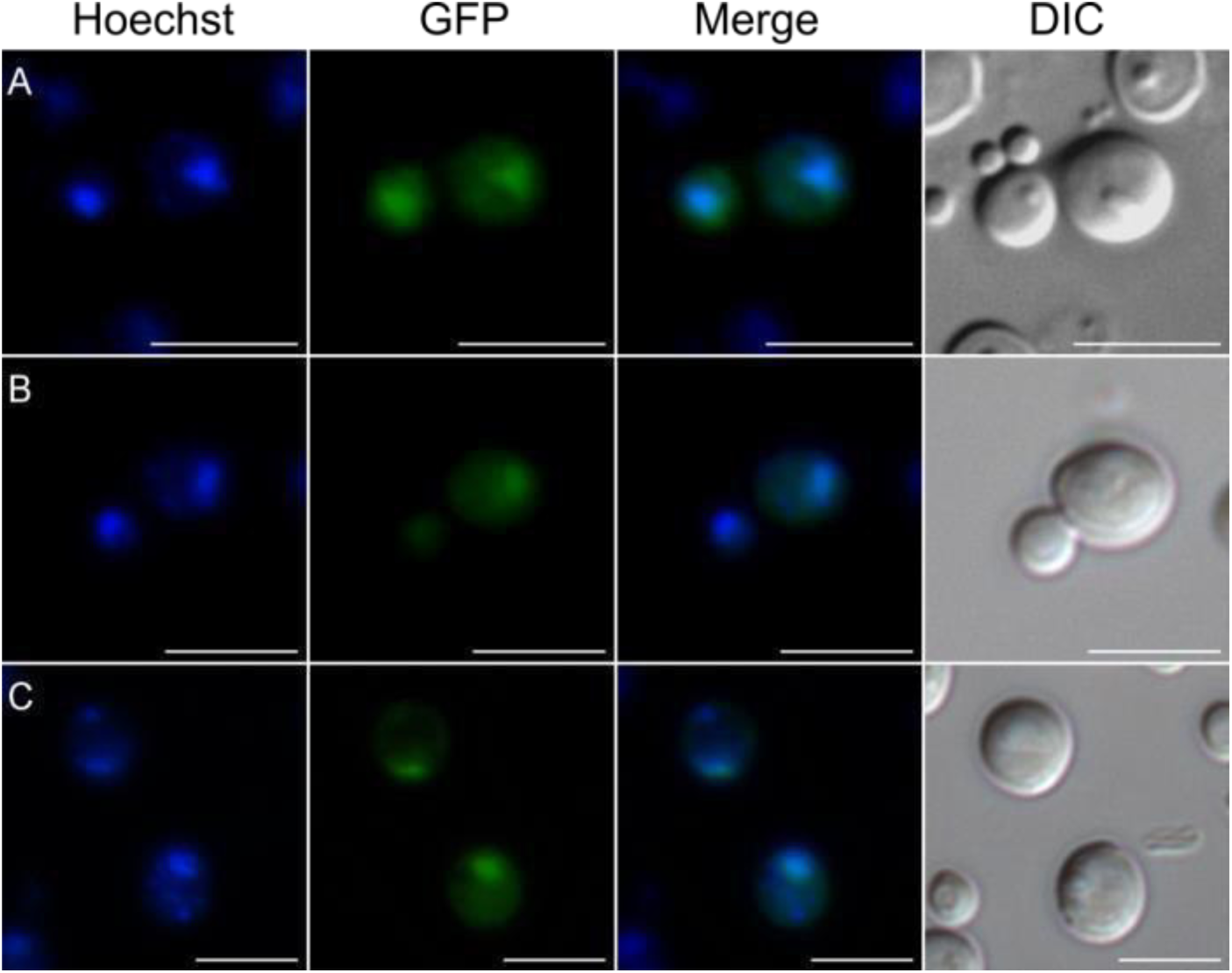
Localization of the SV40-NLS-fused KlGal80p variants and KlGal80-S2p. Localization of c-myc-tagged KlGal80p and its variants was analyzed by immunofluorescence-microscopy. (**A**) The strains JA6/G80M (wild-type KlGal80p), (**B**) JA6/G80-KR56A (KlGal80-K5A/R6Ap), (**C**) JA6/G80-SV40 (wild-type KlGal80p with Nterminal SV40-NLS) and (**D**) JA6/G80-SVKR (KlGal80-K5A/R6Ap with SV40-NLS) were grown in medium containing 2 % galactose. The nucleus was stained with Hoechst 33342. KlGal80 was detected by its c-myc-tag using a primary myc-antibody (Roche) and a secondary Alexa 488 antibody (life technologies). DIC: differential interference contrast. Scale bar: 2 µm. (**E**) The Klgal80 deletion strain JA6/D802R was transformed with pEGFP-G80-S2 (coding for GFP-fused KlGal80-S2p). The cells were grown in medium containing 2 % glucose. Scale bar: 5 µm.

**Supplementary Figure S3:**
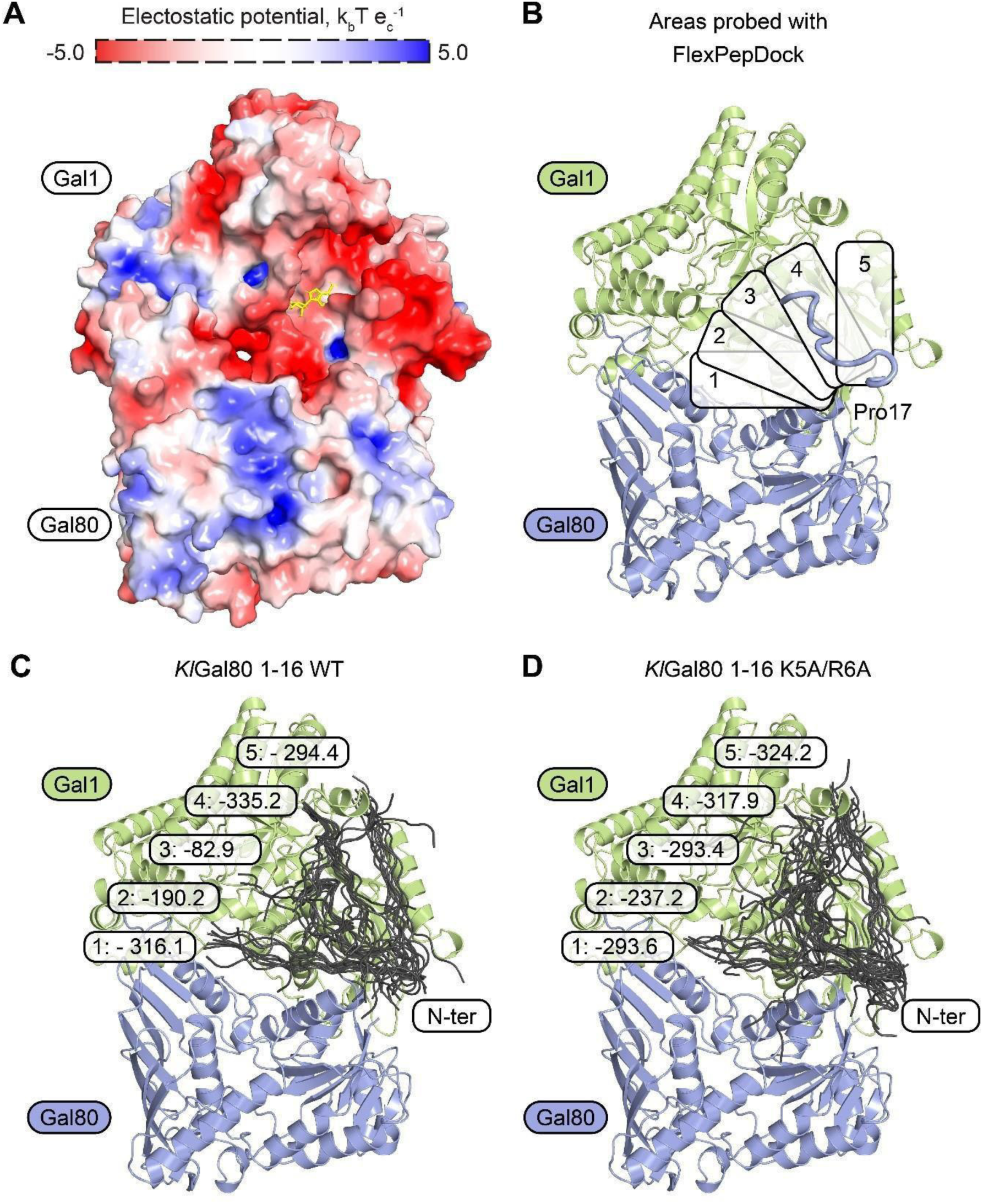
Localization of the GFP-ScGal80-S8K protein. Localization of GFP fused ScGal80-S8Kp was analyzed by fluorescence-microscopy. The plasmid pEGFPScG80-S8K coding for GFP fused ScGal80p with amino acid exchange S8K was transformed into the *gal80* deletion strain JA6/D802R. Cells were grown in minimal medium with (**A**) 2 % glucose, (**B**) 2 % galactose or (**C**) 3 % glycerol and analyzed by fluorescence microscopy. The nucleus was stained by Hoechst 33342. DIC: differential interference contrast. Scale bar: 5 µm.

**Figure S4:**
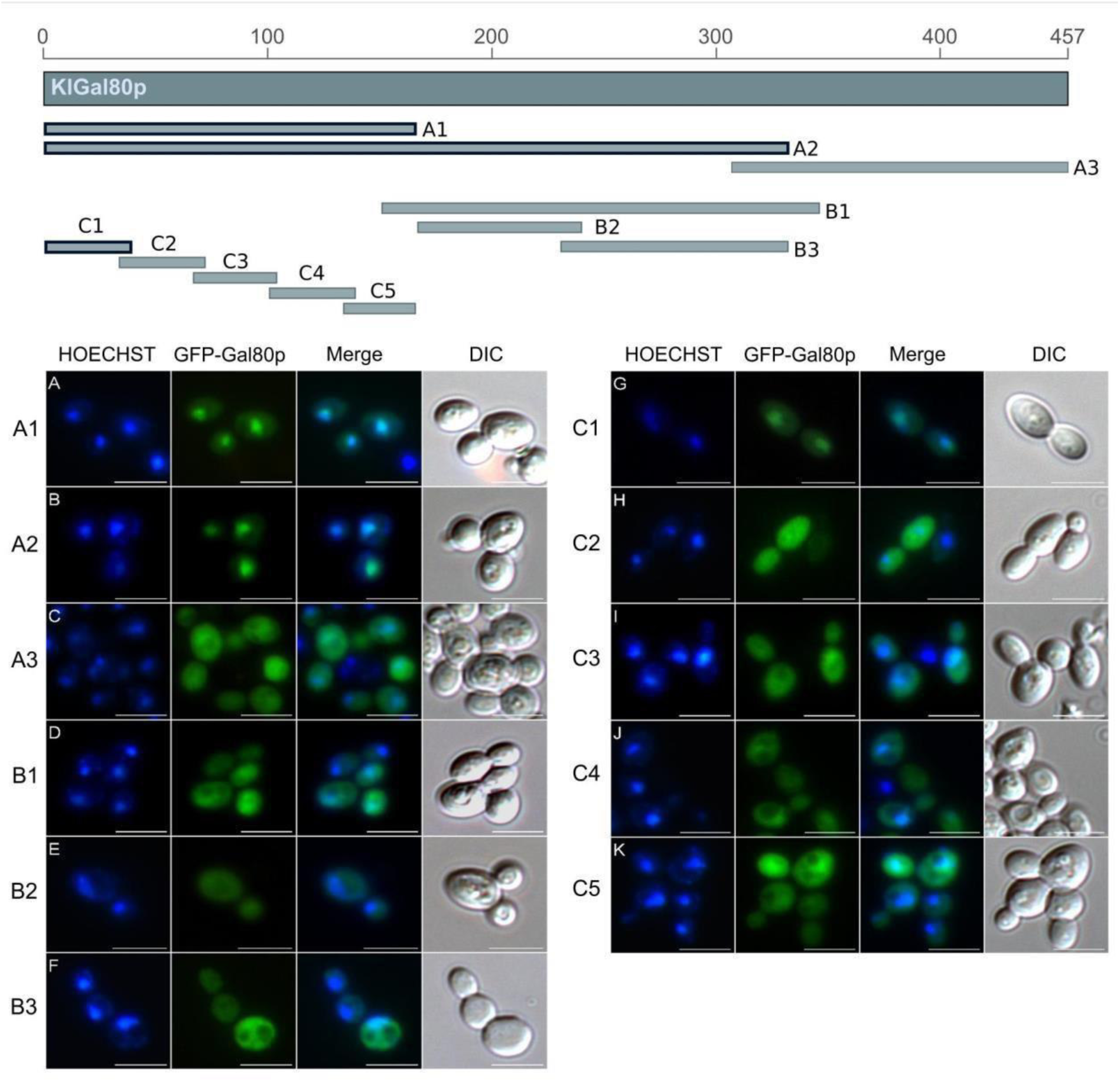
Docking of the KlGal80p N-terminal peptide with KlGal1p. (**A**) The electrostatic charge of a modelled *Kl*Gal1-Gal80 complex. ATP molecule (yellow) is placed based on PDB ID 3V2U. (**B**) Five different areas on the *Kl*Gal1 protein surface were probed for interaction with *Kl*Gal80 1-16 peptide using a FlexPepDock platform. The top scored result is shown. (**C, D**) Structural overview of docking results. In each area the best 10 fits for *Kl*Gal80 1-16 WT (**C**) and K5A/R6A (**D**) are shown. Numbers represent the lowest Rosetta score (lower indicates higher binding probability) in a particular area.

**Supplementary Table S1:** Overview of yeast strains

| Strain | Genotype | Species | Origin |
| --- | --- | --- | --- |
| JA6/G80-KR56A | *gal80-K5A, R6A* | *K. lactis* | This work |
| JA6/G80M | *GAL80* | *K. lactis* | This work |
| JA6/G80-SV40 | *SV40-NLS-GAL80* | *K. lactis* | This work |
| JA6/G80-SVKR | *SV40-NLS-gal80-K5A, R6A* | *K. lactis* | This work |
| FI4 sin4Δ ScGAL80 | *ScGAL80* | *S. cerevisiae* | This work |
| FI4 sin4Δ<br>ScGAL80KR56A | *Scgal80-K5A, R6A* | *S. cerevisiae* | This work |

**Supplementary Table S2:**
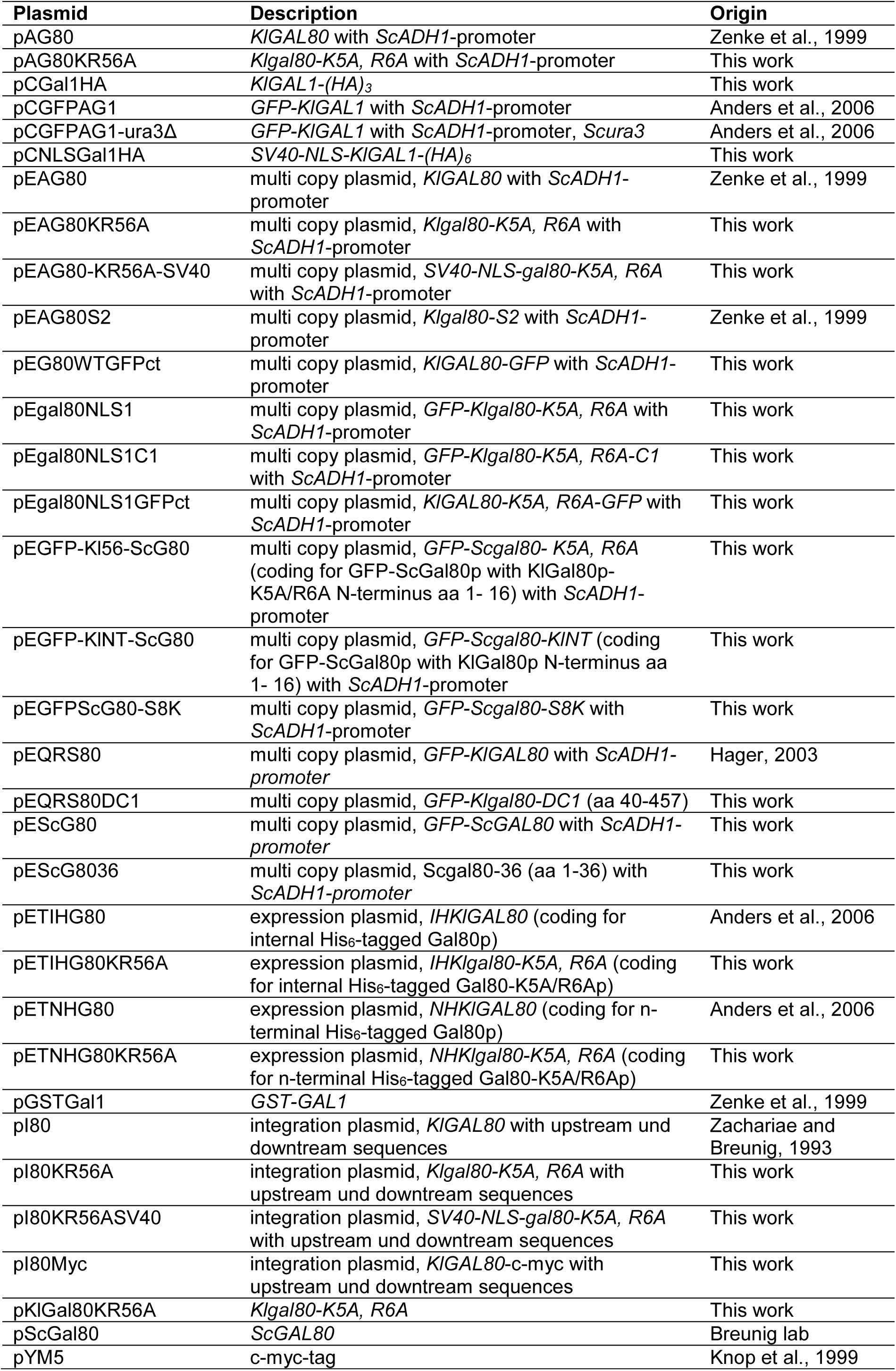
**Overview of used** plasmids

**Supplementary Table S3:**
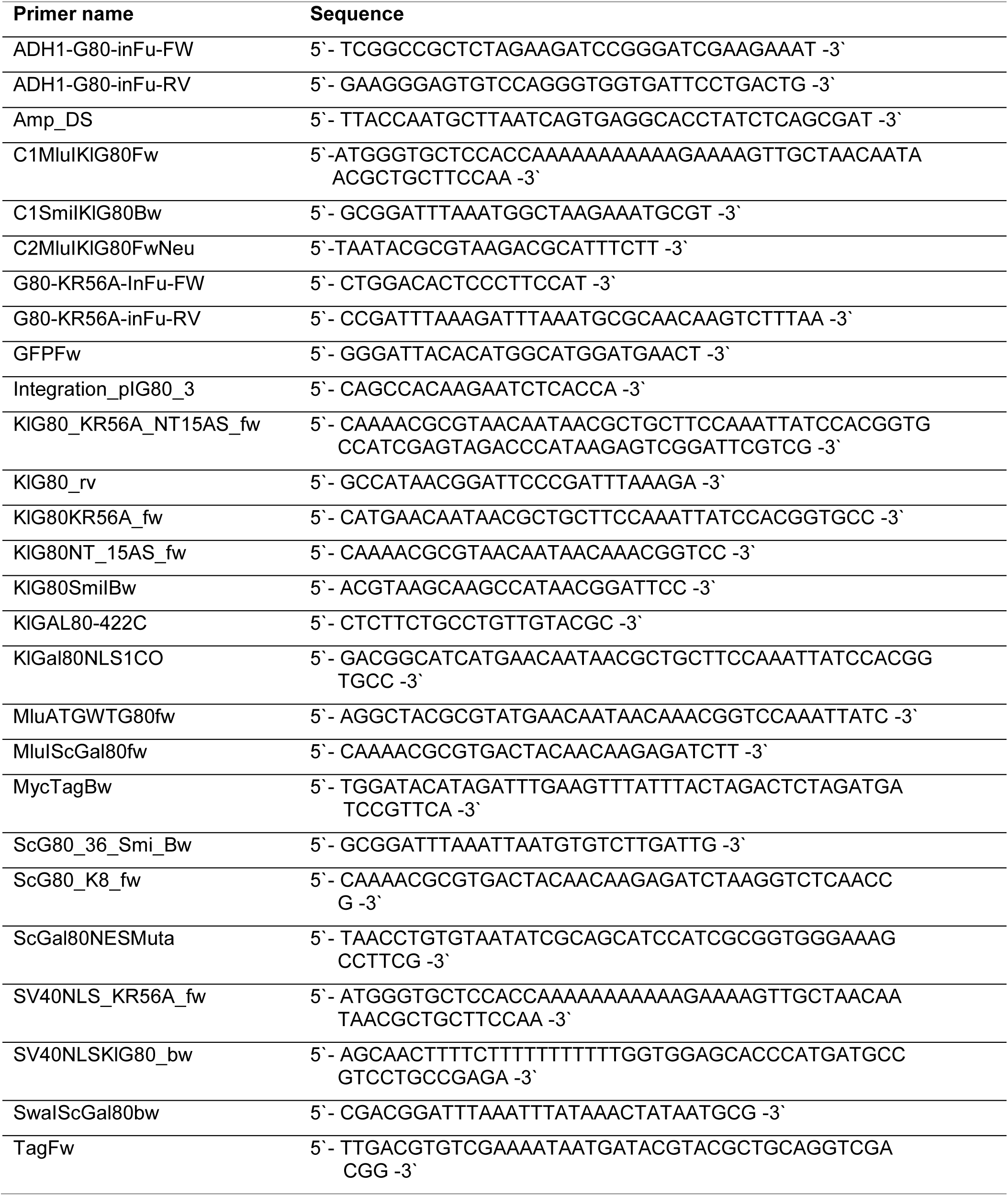
**Overview of DNA** oligonucleotide **sequences used for cloning**

**Supplementary Table S4:**
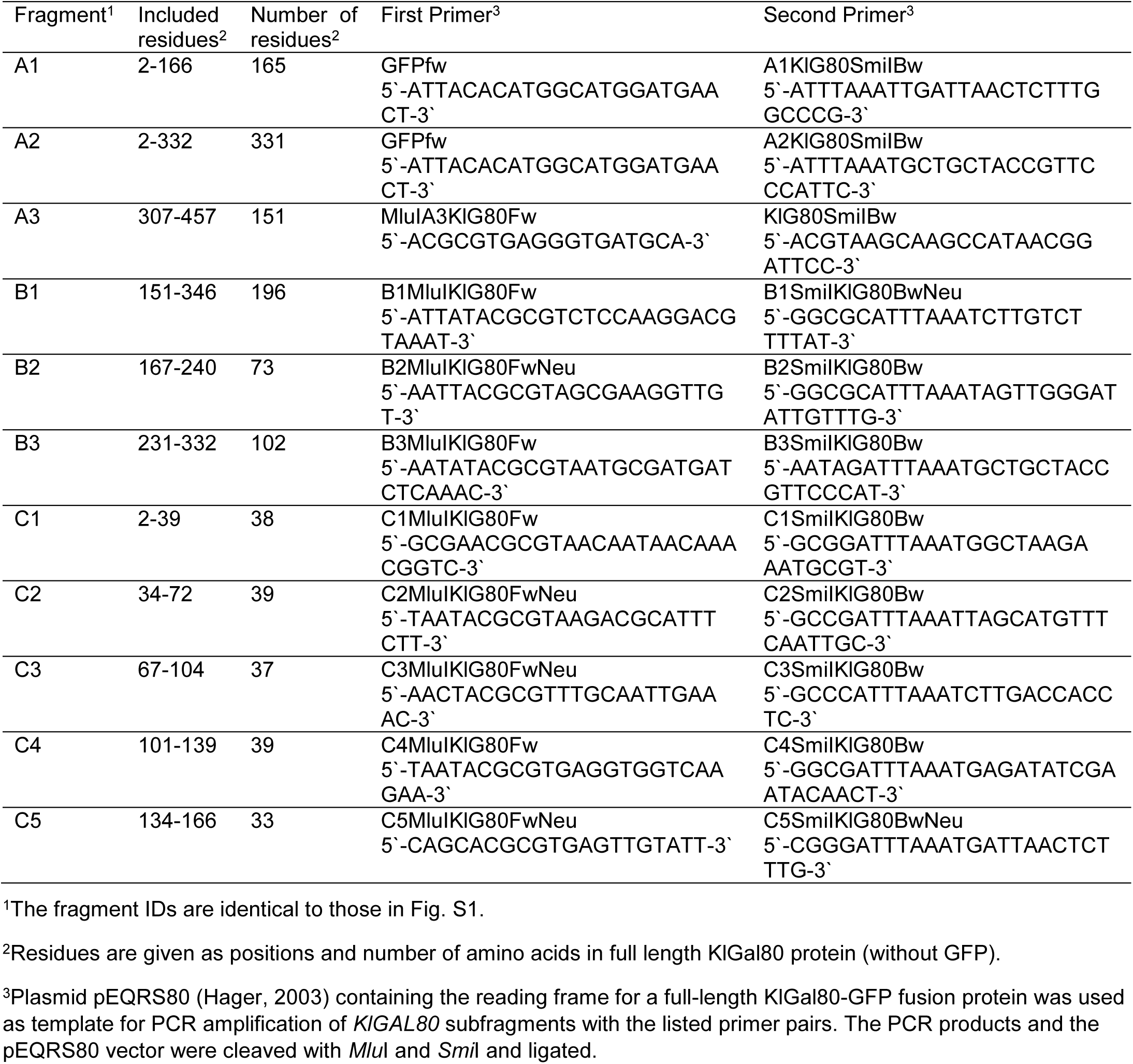
**Overview of** oligonucleotides **used for KlGal80 fragment construction**

